# The salivary protein Saglin facilitates efficient midgut colonization of *Anopheles* mosquitoes by malaria parasites

**DOI:** 10.1101/2022.04.25.489337

**Authors:** Dennis Klug, Amandine Gautier, Eric Calvo, Eric Marois, Stéphanie A Blandin

## Abstract

Malaria is caused by the unicelullar parasite *Plasmodium* which is transmitted to humans through the bite of infected female *Anopheles* mosquitoes. To initiate sexual reproduction and to infect the midgut of the mosquito, *Plasmodium* gametocytes are able to recognize the intestinal environment after being ingested during blood feeding. A shift in temperature, pH change and the presence of the insect-specific compound xanthurenic acid have been shown to be important stimuli perceived by gametocytes to become activated and proceed to sexual reproduction. Here we report that the salivary protein Saglin, previously proposed to be a receptor for the recognition of salivary glands by sporozoites, facilitates *Plasmodium* colonization of the mosquito midgut, but does not contribute to salivary gland invasion. In mosquito mutants lacking Saglin, *Plasmodium* infection of *Anopheles* females is greatly reduced, resulting in impaired transmission of sporozoites at low infection densities. Interestingly, our results suggest that Saglin, present in mosquito saliva, must be ingested with the blood meal to affect *Plasmodium* development, indicating a previously unknown host-pathogen interaction. Furthermore, we were able to show that *saglin* deletion has no fitness cost in laboratory conditions, suggesting this gene would be an interesting target for gene drive approaches.

## Introduction

Vector borne diseases can be caused by a broad range of different pathogens including viruses (e.g. Dengue, DENV), bacteria (e.g. *Borrelia*) and unicellular eukaryotes (e.g. *Plasmodium*). Most of these pathogens have in common that successful transmission requires the infection of the salivary glands of a blood feeding vector, often a tick or a mosquito, to be transmitted with the inoculation of saliva during a blood meal. Still, different pathogens pursue different strategies to achieve this colonization. While vector borne viruses like DENV cause a systemic infection of the mosquito including the salivary glands (Salazar *et al*., 2007), the eukaryotic parasite *Plasmodium* is able to specifically recognize and invade the gland tissue. *Plasmodium* parasites are the causative agent of malaria, the most devastating vector borne disease being responsible for more than half a million fatalities each year (WHO, 2021). *Plasmodium* parasites colonize the midgut of female *Anopheles* mosquitoes following a blood meal from an infected host. Once ingested, *Plasmodium* gametocytes sense the midgut environment through the change in pH, the drop in temperature and the presence of the insect-specific molecule xanthurenic acid (Billker *et al*., 1998) and shed their red blood cell membrane upon activation. Subsequently male gametocytes rapidly divide into eight microgametes which engage in active motility to find and fuse with female macrogametes. After gamete fusion a zygote is formed and differentiates within 20-24 hours into a motile parasite stage, called ookinete. Ookinetes traverse the midgut epithelium to nest below the basal lamina of the mosquito midgut, facing the hemocoele. In this place ookinetes continue development into oocysts that grow for 10-12 days and subsequently divide into ∼1.500-5.000 half-moon shaped sporozoites (Rosenberg and Rungsiwongse, 1991; Hillyer, Barreau and Vernick, 2007). This highly motile parasite stage is released from the oocyst into the hemolymph from where the sporozoites invade the salivary glands to be transmitted to a naive host with the next blood meal. Although the precise kinetic of sporozoite invasion into the salivary glands is unknown, it is believed that the time from release into the hemolymph to successful invasion into the salivary gland is only a few minutes (Douglas *et al*., 2015). Many sporozoites become stuck in the circulatory system of the mosquito (Hillyer, Barreau and Vernick, 2007), and consequently only 10-20% of midgut sporozoites successfully invade the salivary glands (Sinden *et al*., 2007; Mueller *et al*., 2010). Based on the observation that sporozoites invade exclusively gland tissue, it has been hypothesized that sporozoites possess a specific receptor that recognizes a ligand expressed on the outside of the gland. Indeed, *P. knowlesi* is able to infect and invade the salivary glands of *A. dirus* females while it is unable to invade the salivary glands of *A. freeborni*, although midgut colonization in this species proceeds normally. Strikingly, naive *A. dirus* salivary glands transplanted into *P. knowlesi* infected *A. freeborni* females were successfully invaded whereas conversely, no invasion could be observed into glands of *A. freeborni* transplanted in *A. dirus* (Rosenberg, 1985). These observations suggest a limited protein interface that is subject to evolutionary selection pressure so that transmission of *Plasmodium* adapts to the mosquito species that is locally suitable as a vector. Still, the knowledge about salivary gland proteins involved in mediating sporozoite recognition and invasion is sparse, and only few proteins have been characterized in more detail. Based on screens probing the interaction of the two abundant sporozoite surface proteins TRAP and CSP with proteins resident on the salivary glands, Saglin, SG1-like protein and the CSP binding protein (CSP-BP), have been identified as putative ligands for sporozoite entry into salivary glands (Ghosh *et al*., 2009; Wang *et al*., 2013). The SG1-like protein, considered similar to SG1 proteins despite a major difference in size, belongs to a family of proteins with high molecular weight (SGS family) which is conserved across *Anopheles*, *Culex* and *Aedes*. In accordance with these observations a member of the SGS family in *Aedes aegypti*, aaSGS1, was shown to play a role in the salivary gland invasion process of *P. gallinaceum* sporozoites (Korochkina *et al*., 2006). However, a recent study on aaSGS1 using knockout mosquitoes has shown that this protein mainly affects the colonization of the midgut by *P. gallinaceum* rather than the invasion of sporozoites into the salivary glands (Kojin *et al*., 2021). Another interesting protein potentially involved in sporozoite invasion into the salivary glands is CSP-BP, a one to one orthologue of *Drosophila* UPF3. UPF3 belongs to a family of RNA binding proteins which are implicated in mediating quality control of mRNAs (Karousis, Nasif and Mühlemann, 2016; Kojin and Adelman, 2019). How UPF3, which is believed to function intracellularly, mediates sporozoite invasion still needs to be investigated. By far the best characterized salivary gland protein thought to mediate *Plasmodium* sporozoite recognition and invasion is Saglin. Saglin was identified by mass spectrometry as a component of *Anopheles* saliva and has been biochemically characterized as a 50 kDa protein forming a disulphide bonded homodimer (Okulate *et al*., 2007). Interestingly, ingestion of a monoclonal antibody targeting Saglin inhibited the salivary gland invasion rate of *P. yoelii* sporozoites by 73% (Brennan *et al*., 2000) while intrathoracic injection of the same antibody blocked invasion of *P. falciparum* sporozoites by 95% (Ghosh *et al*., 2009). The circular peptide SM1 interacts with both the sporozoite surface protein TRAP and Saglin, and the presence of SM1 blocks gland colonization by sporozoites (Ghosh, Ribolla and Jacobs-Lorena, 2001), leading to the model that recognition of the salivary gland by TRAP is a prerequisite for sporozoite entry (Ghosh *et al*., 2009). Accordingly, the RNAi mediated knockdown of Saglin has also been shown to reduce the efficiency of sporozoite penetration into the salivary glands (Ghosh *et al*., 2009). However, artificial overexpression of Saglin in the distal lobes of the salivary gland failed to boost sporozoite loads, raising doubts about the function of Saglin in sporozoite invasion (O’Brochta *et al*., 2019). Saglin belongs to the SG1 family composed of small salivary gland proteins with unknown functions. In *Anopheles gambiae,* the SG1 family consists of seven members (**Fig. 1**), all of which are specifically expressed in females (Arcà *et al*., 2005). In line with this observation, most SG1 genes are encoded on the X-chromosome, while only one member, *AgTRIO*, is found on an autosome (2R) (**Fig. 1**). Recently the expression pattern of *trio* and *saglin* promoters has been investigated by creating reporter lines expressing fluorescent proteins (Klug *et al*., 2021). Both *trio* and *saglin* promoters drive expression exclusively in the median lobe of the female salivary gland, but they display different activation pattern (**Fig. 1**). In addition, the intergenic 5’ sequence upstream of the *saglin* gene appears to lack intrinsic transcriptional activity, suggesting that the closely clustered X-linked SG1 genes likely share common transcriptional elements indicative of a similar expression pattern (Klug *et al*., 2021). Beside Saglin, the SG1 protein AgTRIO has also been shown to affect sporozoite behaviour. Indeed, knock-down of *AgTRIO* in *P. berghei* infected mosquitoes negatively affects sporozoite transmission (Chuang *et al*., 2019), and administration of α-AgTRIO antibodies to mice significantly reduced infectivity of sporozoites in these mice (Dragovic *et al*., 2018). Still, the molecular mechanisms underlying SG1 protein effect on sporozoite biology remain largely unknown. This study aims at better understanding how Saglin affects *Plasmodium* development in the mosquito.

**Figure 1.**
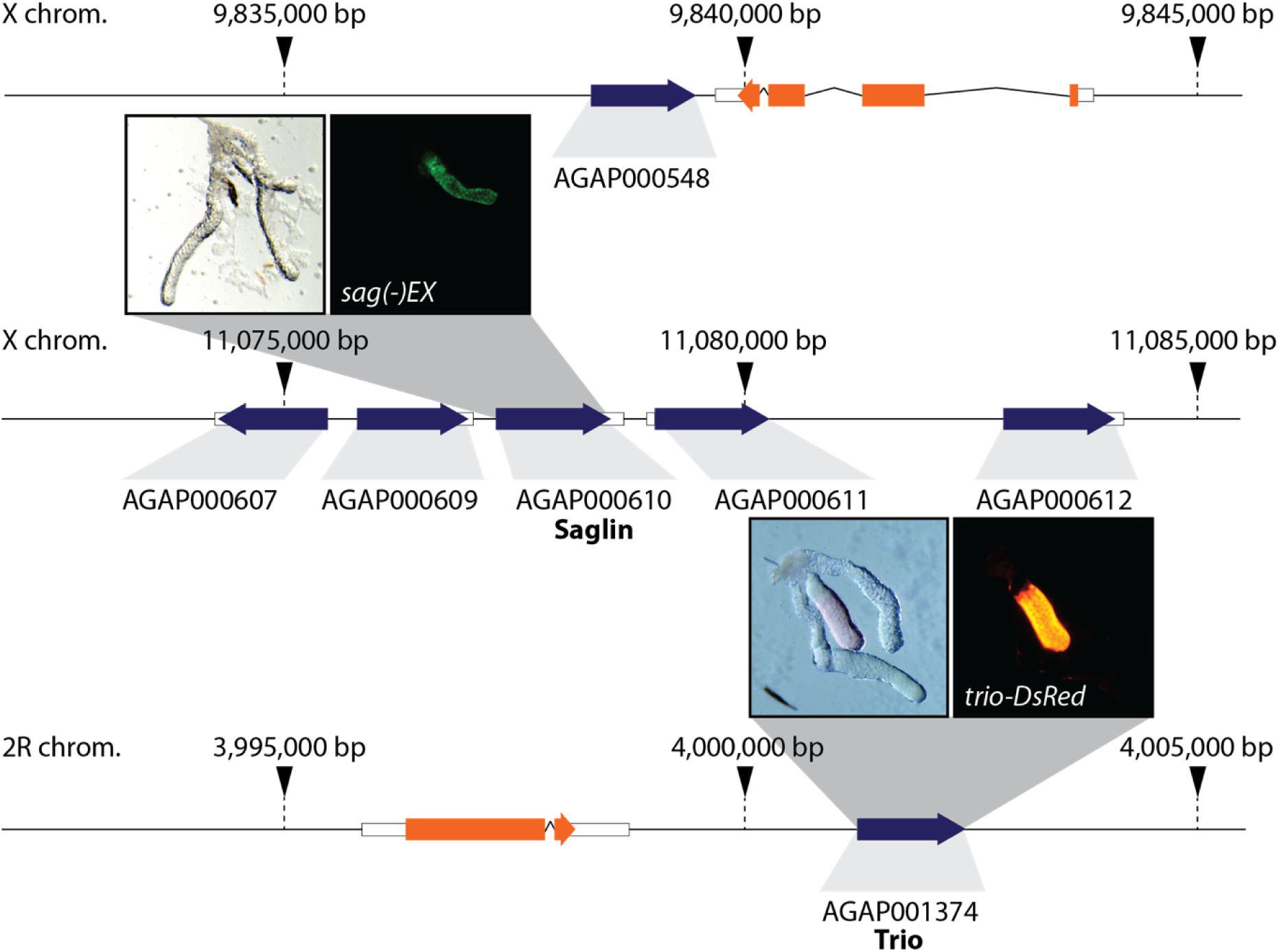
SG1 family proteins in the *Anopheles gambiae* complex. Genomic organization of the SG1 family according to VectorBase (Release 54). Genes encoding SG1 proteins are shown in blue and unrelated genes in orange. Note the high proportion of x-linked genes and the absence of introns in the entire protein family. Representative images of salivary glands dissected from a *sag(-)EX* and a *trio-DsRed* female where EGFP and DsRed are expressed under the control of the *saglin* and *trio* promoters. Both promoters are active in the median lobe.

## Key Resources Table

| Key Resources Table |  |  |  |  |
| --- | --- | --- | --- | --- |
| Reagent type (species) or resource | Designation | Source or reference | Identifiers | Additional information |
| Strain, (*Anopheles coluzzii*) | *Ngousso* | Morlais Isabelle, IRD/OCEAC, Yaounde | MRA-1279 | Wild-type strain |
| Strain, (*Anopheles coluzzii*) | *aapp-DsRed* | (Klug *et al.*, 2021) | / | DsRed expression in distal lobes of the salivary gland |
| Strain, (*Anopheles coluzzii*) | *trio-DsRed* | (Klug *et al.*, 2021) | / | DsRed expression in median lobe of the salivary gland |
| Strain, (*Anopheles coluzzii*) | *sag(-)KI* | (Klug *et al.*, 2021) | / | Knockin into *saglin* gene expressing EGFP from 3xP3 and endogenous *Saglin* promoters |
| Strain, (*Anopheles coluzzii*) | *sag(-)EX* | (Klug *et al.*, 2021) | / | Knockin into *saglin* with excised 3xP3 promoter expressing EGFP from endogenous *saglin* promoter |
| Strain, (*Mus musculus*) | *CD-1* | Janvier Labs, Le Genest-Saint-Isle, France | / | self-bred in in-house facility |
| Strain,<br>(*Plasmodium berghei*) | $\Delta p230p$-<br>GFP | (Manzoni *et al.*,<br>2014) | / | Constitutive<br>expression of<br>GFP along<br>the life cycle |
| Strain,<br>(*Plasmodium falciparum*) | NF54 | (Bryant *et al.*,<br>2018) | / | / |
| Protein | recombinant<br>Saglin | this study | / | / |
| Antibody | α-PPO2 | (Fraiture *et al.*,<br>2009) | / | Rabbit<br>polyclonal |
| Antibody | α-Saglin | this study | / | Rabbit<br>polyclonal |
| Antibody | α-GFP | ThermoFisher<br>Scientific,<br>Waltham, US | Cat#PA1-980A | Rabbit<br>polyclonal |
| Antibody | α-mCherry | abcam,<br>Cambridge, UK | Cat#167453 | Rabbit<br>polyclonal |
| Antibody | α-TEP1 | (Blandin *et al.*,<br>2004) | / | Rabbit<br>polyclonal |
| Antibody | Antibody coupled to HRP | Promega, Madison, US | Cat#W4011 | Goat anti rabbit secondary antibody |
| Antibody | Antibody coupled to Alexa Fluor 546 | ThermoFisher Scientific, Waltham, US | Cat#A-11035 | Goat anti rabbit secondary antibody |
| Software, algorithm | Prism 5.0 | GraphPad, San Diego, US | / | https://www.graphpad.com/scientific-software/prism/ |
| Software, algorithm | FlowJo v10.8.1. | BD Biosciences, Ashland, US | / | https://www.flowjo.com/ |
| Software, algorithm | FIJI | (Schindelin *et al.*, 2012) | / | https://imagej.net/software/fiji/ |
| Software, algorithm | MetaMorph | Molecular Devices, San Jose, US | / | https://www.moleculardevices.com/contact |
| Software, algorithm | VectorBase (version 36-55) | (Amos *et al.*, 2022) | / | https://vectorbase.org/vectorbase/app |
| Software, algorithm | SignalP 5.0 | (Almagro Armenteros *et al.*, 2019) | / | https://services.healthtech.dtu.dk/service.php?SignalP-5.0 |
| Software, algorithm | SMART | (Letunic, Khedkar and Bork, 2021) | / | http://smart.embl-heidelberg.de/ |
| Software, algorithm | NIS-Elements, F 4.20.01, 64 Bit | Nikon Instruments Inc., Melville, US | / | https://www.microscope.healthcare.nikon.com/products/software/nis-elements |
| Other | WGA-XFD594 | AAT Bioquest, Sunnyvale, US | Cat#25509 | / |
| Other | Hoechst 33342 | ThermoFisher Scientific, Waltham, US | Cat#H3570 | / |
| Other | Ponceau S solution | Sigma-Aldrich, St Louis, US | Cat#P7170-1L | / |
| Other | Diff-Quik Solution I and II | RAL Diagnostics, Martillac, France | Cat#10736133<br>Cat#10736134 | / |

## Material & Methods

### Animals

Knock-in transgenesis was performed in a *vasa-Cas9* expressing *A. coluzzii* strain which had been introgressed into the *Ngousso* genetic background (Keller Valsecchi *et al*., 2021). As a positive control of GFP expression in the midgut we used the *A. stephensi* line G12-GFP (Nolan *et al*., 2011). Male CD-1 mice (purchased from Janvier Labs and further self-bred at the in-house facility) were used to perform *P. berghei* infections and female CD-1 mice were used for routine blood feedings to propagate mosquito colonies.

### Mosquito breeding

*Anopheles coluzzii* and *Anopheles stephensi* mosquitoes were reared in standard conditions (27°C, 75-80% humidity, 12-hr/12-hr light/dark cycle) as described previously (Volohonsky *et al*., 2017). In brief, larvae were reared in osmosed water and fed daily pulverized fish food (TetraMin). Pupae were collected in small glass dishes and transferred into netted cages. Hatched mosquitoes were fed on 10% sugar solution *ad libitum*. To propagate colonies, ≥4 day old mosquitoes were blood fed for 10-15 minutes on anesthetized mice. Two-three days later a glass dish with wet filter paper was provided to allow egg laying. Larvae hatched approximately 48 hours after the deposition of eggs. Larvae were floated in demineralized water and reared as described.

### Estimating fitness of the *sag(-)KI* transgene by flow cytometry

A Complex Object Parametric Analyzer and Sorter (COPAS, Union Biometrica) was used as described (Marois *et al*., 2012; Bernardini *et al*., 2014) to quantify the frequency of the *sag(-)KI* transgene in mixed populations of neonate mosquito larvae and to count and mix defined numbers of homozygous *sag(-)KI* and wild-type larvae for experiments. Flow cytometry was always performed on unfed neonatal larvae in water free of any debris to minimise background fluorescence and ensure accurate sorting. To assess the fitness of the *sag(-)KI* transgene, a mosquito colony was started by crossing 24 homozygous *sag(-)KI* females with 24 wild-type males. Subsequently the frequency of *sag(-)KI* carriers and non-carriers was measured by COPAS sorting of neonate larvae for 10 generations. Flow cytometric data were analyzed using FlowJo v10.8.1. and ratios of transgene carriers and non-carriers were plotted using GraphPad Prism 5.0.

### Infections with *Plasmodium berghei*

To infect mosquitoes with the rodent malaria parasite *P. berghei,* male CD1 mice were injected intraperitoneally with frozen parasite stocks. Infections were performed with the *P. berghei* line Δp230p-GFP (Manzoni et al., 2014) genetically engineered to constitutively express EGFP under the control of the promoter of heat shock protein 70 (*hsp70*). Upon infection parasitemia was monitored by flow cytometry (AccuriC6 SORP, Becton Dickinson). Once parasitemia reached ≥1%, mice were bled by cardiac puncture and blood stage parasites were transferred into naïve mice by intravenous injection into the tail vein. Usually 2.10^7^ parasites were transferred per mouse to achieve a parasitemia of 1.5-3.5% three days post injection, when mosquitoes were allowed to feed on the anaesthetized mice (same conditions as for regular blood feeding). Parasites were cycled regularly through mosquitoes to maintain fitness. Parasite stocks were stored at -80°C by mixing equal volumes of infected blood with a parasitemia of ≥1% and phosphate buffered saline (PBS) supplemented with 30% glycerol.

### Culture of *P. falciparum* and mosquito infection

Gametocytes were produced *in vitro* using a *Plasmodium falciparum* NF54 clone (Bryant *et al*., 2018) and similar procedures as in (Tripathi *et al*., 2020). All cultures were maintained in A+ human red blood cells (EFS, Strasbourg) at 3.8% haematocrit and complete medium (RPMI 1640 with L-glutamine and 25mM HEPES supplemented with 10% human A+ serum (EFS, Strasbourg) and 10 mM hypoxanthine (c-c-Pro, Oberdorla)), at 37°C, under a 5% O2, 5% CO2 and 90% N2 atmosphere. Briefly, asexual parasite cultures were kept at 3% parasitemia maximum. Gametocyte cultures were seeded in a 6-well plate at 0.5% parasitemia and maintained in culture with daily medium changes for 16 days before infection. The quality and density of gametocytes was regularly evaluated on blood smears. Gametocyte infected RBCs were mixed with non-infected RBCs and human serum (50% hematocrit) and offered to starved female mosquitoes using a Hemotek system mounted with parafilm to maintain the blood meal at 37°C. Unfed females were discarded 1-3h post feeding, and the remaining maintained at 27°C, 70% humidity for 7-8 days before dissection.

### Parasite counting in mosquito midguts and salivary glands

To assess parasite burdens, infected mosquitoes were anesthetized on ice and transferred into a Petri dish containing 70% ethanol. After 1-2 minutes mosquitoes were transferred into a second Petri dish containing phosphate buffered saline (PBS). For oocyst counting, mosquitoes were dissected 7-8 days post infection (dpi) on a microscope slide in a drop of PBS and a SZM-2 Zoom Trinocular stereomicroscope (Optika). Dissected midguts infected with GFP-expressing *P. berghei* were transferred on a second microscope slide in a drop of fresh PBS, covered with a cover slip and imaged using a Nikon SMZ18 fluorescence stereomicroscope. Images were then processed using the watershed segmentation plugin (Soille and Vincent, 1990) and oocysts were subsequently counted using the „analyze particles“ function implemented in Fiji (Schindelin *et al*., 2012). Midguts infected with *P. falciparum* were dissected in PBS and stained with 0.2% mercurochrome in water. Samples were washed three times in PBS for 10 min each before imaging using a Zeiss Axio Zoom.v16 microscope. *P. falciparum* oocysts were counted manually using Fiji (Schindelin *et al*., 2012).

For sporozoite counting, pools of salivary glands from ≥10 mosquitoes (17-18 dpi) were dissected and manually grinded in 50 µl PBS with a plastic pestle for one minute in a 1,5 ml plastic reaction tube. Samples were diluted by adding 50 µl of PBS used to carefully rinse the pestle. Subsequently 7 µl of sample solution were loaded into a Neubauer hemocytometer. Sporozoites were allowed to settle for 5 minutes and counted using a light microscope (Leica DFC3000 G) with 20-fold magnification.

### Analysis of transmission capability *in vivo*

Freshly hatched *sag(-)KI* larvae obtained from a homozygous colony were mixed in equal amounts with wild-type larvae (*Ngousso*), raised together and infected with the GFP-expressing *P. berghei* as described above. Blood fed females were collected and kept at 21°C and 60-70% humidity. Seven days after blood feeding, midguts from a random sample of mosquitoes were dissected to monitor parasite load and prevalence. On the 15th day after infection mosquitoes were briefly anaesthetized on ice and sorted as wild-type or *sag(-)KI* females according to their EGFP expression in the eye (typical expression pattern of the 3xP3 promoter). Subsequently groups of 10 mosquitoes of identical genotype were distributed in paper cups. On the evening of the 17th day after infection sugar pads on cups were removed to starve mosquitoes overnight. The following day one anaesthetized mouse was placed on each cup and mosquitoes were allowed to blood feed for 15 minutes. A minimum of three and a maximum of eight mosquitoes per cup were observed to have successfully taken blood. Parasitemia in mice was monitored by flow cytometry (AccuriC6 SORP, Becton Dickinson) from day three to day seven after biting. On day seven, all infected animals were sacrificed to avoid suffering and non-infected mice followed for an additional week. All transmission experiments were performed with male CD1 mice with an approximate age of six months.

### Analysis of blood feeding behaviour

Neonate homozygous *sag(-)KI* and wild-type larvae of the same colony were either COPAS sorted or obtained from homozygous *sag(-)KI* and control *Ngousso* colonies, and raised as a mix. Adult females aged ≥4 days were allowed to blood feed for 10 minutes. After blood feeding females were anaesthetized on ice and sorted to discard unfed females and, when necessary, to count GFP-positive and GFP-negative individuals to calculate the feeding efficiency for both genotypes. Blood fed females were shock frozen for 10 minutes at -20°C and subsequently weighed using a fine scale (Ohaus, Explorer Pro). For the calculation of the number of ingested cells, blood fed females were kept unfrozen. Their abdomen was dissected, transferred into a plastic reaction tube containing either 1500 µl or 2000 µl PBS and crushed with a pestle to release the ingested blood. Seven µl of the solution was loaded in a Neubauer counting chamber. Red blood cells were allowed to settle for 5 min before counting.

### Exflagellation assay

Four to ten-day old *sag(-)KI* and wild-type mosquitoes were allowed to take a blood meal on an infected mouse for five minutes. Mosquitoes were then kept at 21°C and 60-70% humidity for 12 minutes to activate gametocytes and induce exflagellation of male gametes, briefly anaesthetized on ice and their midguts were opened to release and smear the blood meal on a microcopy slide using two syringes. Blood smears were air dried, fixed for one minute in methanol and stained using Diff-Quik (RAL Diagnostics). Male gametes were counted using a light microscope (Leica DFC3000 G) with 100-fold magnification and a counting grid. For each sample, the microgametes in 30 fields and the number of red blood cells in one representative field were counted to calculate the microgametocytemia.

### Fluorescence imaging

Imaging of DsRed or EGFP in mosquitoes, midguts and salivary glands was performed using a Nikon SMZ18 stereomicroscope with a Lumencor Sola Light engine, in brightfield and with appropriate fluorescence filters. Scale bars were implemented in reference to an objective micrometer (Edmund optics) that was imaged with the same magnification. Images were edited using Fiji (Schindelin *et al*., 2012).

### Expression of recombinant Saglin

The *saglin* gene (AGAP000610, coding for mature peptide only) was codon-optimized for *E. coli* expression and subcloned into pET19b by Biobasic Inc., Markham, Canada. The synthetic gene was designed to contain *NdeI* (5’-end) and *XhoI* (3’end) restriction sites and a 6xHis-tag before the stop codon. Expression of recombinant Saglin in *E. coli* (BL21pLYS) was performed under standard conditions as described previously (Williams *et al*., 2021). Purification of recombinant Saglin was carried out by Immobilized metal affinity chromatography (IMAC) followed by Size-exclusion chromatography (SEC) using the Akta Purifier system (GE Healthcare, Piscataway, NJ). Briefly, cell lysates containing the recombinant protein were supplemented with 500 mM NaCl and 5 mM imidazole and loaded onto a HiTrap Chelating HP column charged with NiCl2 (5 mL bed volume, GE Healthcare, Piscataway, NJ, USA). Fractions were eluted using a linear gradient 0-1 M Imidazole in 25 mM phosphate buffer, 500 mM NaCl, pH 7.4 over 60 min at 2 ml/min. Fractions containing recombinant Saglin were pooled and concentrated using Amicon® Ultra-15 centrifugal filter units (Millipore Sigma) and fractionated on a Superdex 200 10/300 GL column (GE Healthcare, Piscataway, NJ). Purity was verified by NuPAGE 4-12%, Bis-tris (ThermoFisher Scientific), visualized by Coomassie staining. Protein identity was confirmed by N-terminal sequencing using automated Edman degradation (Research Technology Branch, NIAID).

### Generation of α-Saglin antibodies

Polyclonal antibodies against recombinant Saglin were raised in rabbits as described previously (Martin-Martin *et al*., 2020). Briefly, immunization of rabbits (New Zealand White) was carried out in Noble Life Science facility (Woodbine, MD) according to their standard protocol. Rabbits received a total of 3 immunizations with 1 µg of recombinant protein each at days 0, 21, 42. Freund’s Complete Adjuvant was used for the initial immunization and Freund’s Incomplete Adjuvant was utilized for the subsequent boosters. Rabbit sera were collected by exsanguination 2 weeks after the last boost.

### Western blotting of different mosquito tissues

Ten Salivary glands (equivalent of five mosquitoes) or five midguts were dissected in 20 µl Laemmli buffer. Hemolymph collection of 20 female mosquitoes was performed by proboscis clipping directly into 10 µl Laemmli buffer. To collect probosci, mosquitoes were anaesthetized on ice, 20 probosci (equivalent of 20 mosquitoes) were cut using scissors and transferred into 20 µl Laemmli buffer using forceps. To assess distribution of Saglin in the mosquito body, four female mosquitoes with salivary glands removed were collected in 100 µl Laemmli buffer. Salivary glands, midguts, probosci and carcasses were crushed manually using pestles for 30-60 sec before samples were denatured at 65 °C for 5 minutes in a heat block. A stock of recombinant Saglin (1mg / ml) was diluted 1:20 in Laemmli buffer and denatured together with other samples as positive control. Subsequently samples were centrifuged for 3 min at 13.000 rpm and a maximum of 20µl per sample was loaded on a Mini-PROTEAN TGX Stain-Free Precast Gel (BioRad) using 6 µL of Thermo Scientific™ PageRuler™ Plus Prestained Protein Ladder as standard. Separation was performed at 170 V using the Mini Trans-Blot^®^ cell system (BioRad). Once separation was complete, gels were blotted on a PVDF membrane (Trans-Blot Turbo Mini 0.2 µm PVDF Transfer Pack; BioRad) using the mid-range program of a Pierce Fast blotter (Thermo Fisher Scientific). The membrane was blocked for one hour in PBST (PBS containing 0.1% Tween) supplemented with 5% fat-free milk powder. Membranes were incubated overnight at 4°C with primary antibody diluted in PBST supplemented with 3% fat-free milk powder (PBST3M). Blots were washed three times for 10 minutes in PBST and incubated for one hour at room temperature in secondary antibody conjugated to horseradish peroxidase (HRP) diluted in PBST3M. After incubation with secondary antibodies, membranes were washed three times for 10 minutes with PBS. Antibody binding was revealed using the Super signal WestPico Plus kit (Thermo Fisher Scientific). After 1-2 minutes of incubation, images were acquired using the Chemidoc software (Biorad). Before incubation with further primary antibodies to visualize additional proteins, membranes were stripped for 20 to 30 minutes in Restore PLUS Western Blot Stripping Buffer (Thermo Fisher Scientific) and washed three times for 10 minutes in PBST followed by a new incubation in blocking solution.

### Western blotting and Coomassie staining of mosquito midgut contents

Mosquitoes were allowed to feed blood on an anaesthetized BALB/c mouse for a maximum of 5 minutes. Subsequently mosquitoes were anaesthetized on ice and midguts of five blood or sugar-fed siblings were dissected on a microscopy slide. In preparation of the dissection a 20µl drop of PBS premixed with heparin was placed next to each mosquito destined for dissection. Removed midguts were placed within the drop before opening. Subsequently midgut contents of five guts were collected with a pipette and transferred to a fresh plastic reaction tube yielding an approximate volume of 100µl. To avoid degradation of proteins within the blood meal, dissections were performed within 10 minutes after blood feeding. Samples were homogenized by passing through an insulin syringe for 10 times and clarified by centrifugation for 1-2 min at 14.000 g and 4 °C. Subsequently protein concentration was measured (Denovix) and samples were stored at -80°C before use. For electrophoresis 50-75µg of protein per sample were loaded on a Nu-PAGE and separated using MES buffer. Samples were run in duplicate to allow for Coomassie staining and immunoblotting. Membrane transfer was performed using iBLOT 2 (Invitrogen) overnight at 4°C in TBS-T buffer supplemented with 5% milk powder. Immunoblots were incubated with α -Saglin antibodies (1:800) in TBS-T supplemented with 0,05% milk powder, washed four times in TBS-T and incubated with secondary antibody (1:10.000). Immunoblots were developed with the Super signal WestPico Plus kit (Thermo Fisher Scientific).

### Indirect immunofluorescence staining of salivary glands

Immunofluorescence staining of Saglin expression in salivary glands was performed according to (O’Brochta *et al*., 2019). The protocol was modified to treat the samples with PBSBT (PBS supplemented with 1% BSA and 0.1% Triton X-100) only once for 5 minutes and then three times for 10 minutes after treatment with the secondary antibody. The primary α-Saglin antibody and the secondary Alexa Fluor 546 coupled anti-rabbit antibody were both diluted 1:200. Imaging was performed using a Zeiss LSM 780 microscope equipped with a Hamamatsu Orca Flash 4.0 V1 camera using a 63x (NA 1.4) objective.

### Injection of fluorescently labeled Wheat Germ Agglutinin (WGA) in mosquitoes

Mosquitoes 4-10 days post emergence were injected twice using the Nanoject III Injector (Drummond) to reach a total of 100 nl of XFD594 (structural analogue of Alexa Fluor 594) labeled WGA at a concentration of 40 µg/ml. Salivary glands were dissected one hour after injection and imaged with a Zeiss LSM 780 confocal microscope equipped with a Hamamatsu Orca Flash 4.0 V1 camera using a 40x (NA 1.4) objective. Fluorescence patterns were quantified using Fiji (Schindelin *et al*., 2012).

## Data availability

All data have been made available within this manuscript. Transgenic mosquito lines as well as plasmids are available upon request.

## Statistical analysis

Statistical analysis was performed using GraphPad Prism 5.0 (GraphPad, San Diego, CA, USA). Depending on data sets statistics were performed with Mann Whitney test, one-way ANOVA or Fisher’s exact test. A value of p<0.05 was considered significant.

## Ethics statement

Experiments were carried out in conformity with the 2010/63/EU directive of the European Parliament on the protection of animals used for scientific purposes. Our animal care facility received agreement #I-67-482-2 from the veterinary services of the département du Bas Rhin (Direction Départementale de la Protection des Populations). The use of animals for this project, and the generation and use of transgenic lines (bacteria, mosquitoes, parasite) were authorized by the French ministry of higher education, research and innovation under the agreements number APAFIS#20562-2019050313288887 v3, and number 3243, respectively.

## Acknowledgements

We thank Ludivine Ramolu for technical support, Sarra Manai for help with western blot experiments, Lionel Brice Feufack Donfack for help with *P. falciparum* infections and Dr. Jean-Daniel Fauny for assistance during microscopy. We also thank Dr. Paola Valenzuela-Leon and Karina Botello for assistance with midgut Western blots. We would like to thank the mosquito immune responses (MIR) team for fruitful discussions and for assistance with mosquito breeding.

## Authors contributions

DK designed and performed experiments, analyzed data, made figures and wrote the draft. EC provided resources and helped writing the manuscript. EM designed experiments, performed transgenesis of mosquitoes and helped writing the manuscript. SAB provided resources, helped with the experimental design, performed infection experiments and edited the draft. All authors edited and approved the manuscript for publication.

## Results

### Absence of Saglin impairs midgut colonization by *Plasmodium berghei* and *Plasmodium falciparum*

To investigate the impact of the loss of Saglin expression on the susceptibility of mosquitoes to *Plasmodium* infection, we made use of the mosquito line *sag(-)KI* in which the coding sequence of Saglin (AGAP000610) is replaced with EGFP (Klug *et al*., 2021) (**Fig. 2A**). In a first experimental setup, homozygous *sag(-)KI* mosquitoes and wild-type (WT) siblings obtained from the same colony were co-cultured and infected together with the rodent malaria parasite *P. berghei*. Determination of parasite loads in midguts dissected on days 7 and 8 post infection revealed a mean of 31 oocysts in *sag(-)KI* females, whereas WT females were infested with a mean of 122 oocysts indicating a significant difference in the susceptibility to *Plasmodium* infection in *sag(-)KI* females (**Fig. 2B**). To simplify mosquito breeding for subsequent infection experiments, we then bred WT and homozygous *sag(-)KI* mosquitoes as separate colonies and obtained neonate larvae of both colonies derived from a WT and a homozygous *sag(-)KI* colony were either mixed or reared separately, but always synchronously to achieve equal proportions of males and females of both genotypes. To test for possible biases induced by the new breeding scheme, infection experiments were repeated. Infected *sag(-)KI* and WT mosquitoes showed a mean of 29 oocysts while a mean of 56 oocysts was determined for wild-type (*control*) confirming the phenotype (**Fig. 2C**). Despite the drop in oocyst numbers observed in *sag(-)KI* females, prevalence for infection was not significantly different from WT even if prevalence in the *sag(-)KI* population was reproducibly slightly lower compared to the *control* **(Fig. 2B, Fig. 2 – figure supplement 1**). To further test if the observed changes of the parasite burden in the midgut result in lower sporozoite concentrations in the salivary glands, we counted salivary gland sporozoites (SGS) at days 17-18 post infection using a hemocytometer. SGS numbers in *sag(-)KI* females were indeed significantly reduced (**Fig. 2C**). While WT females displayed a mean of 30,400 SGS per mosquito, only 7,100 SGS per mosquito were observed in *sag(-)KI* females (>75% reduction in sporozoite load) (**Fig. 2C**). Based on this observation, we investigated whether the reduction in SGS was solely due to reduced oocyst numbers or whether sporozoites in *sag(-)KI* mosquitoes also exhibit impaired salivary gland recognition, as Saglin has been previously described as a determinant of salivary gland entry (Ghosh *et al*., 2009). We quantified the sporozoites invasion rates in the salivary glands of *sag(-)KI* and WT mosquitoes by calculating the ratio of SGS per oocyst based on results obtained from the same batches of infected mosquitoes. Medians of 175 and 170 SGS per oocyst were obtained for *sag(-)KI* and WT females, respectively, indicating that recognition of the salivary glands by sporozoites in the absence of Saglin is not impaired (**Fig. 2D**). To test whether *sag(-)KI* would also affect mosquito susceptibility to the human malaria parasite *P. falciparum*, we initially infected mosquitoes with a low density of parasites to simulate conditions in the wild (Bompard *et al*., 2020). While 65% of WT mosquitoes carried parasites, only 13% of *Sag(-)KI* mutants were infected, and those infected carried fewer parasites than WT (median of 2 vs 7, respectively) (**Fig. 2E**). The reduced susceptibility of *Sag(-)KI* mosquitoes to *P. falciparum* was further confirmed in a second experiment where mosquitoes were infected with a higher density of *P. falciparum* gametocyte: prevalence of 76% vs 92%, and median load of 11 vs 35 oocysts per midgut in *Sag(-)KI* vs WT, respectively (**Fig. 2F**).

**Figure 2.**
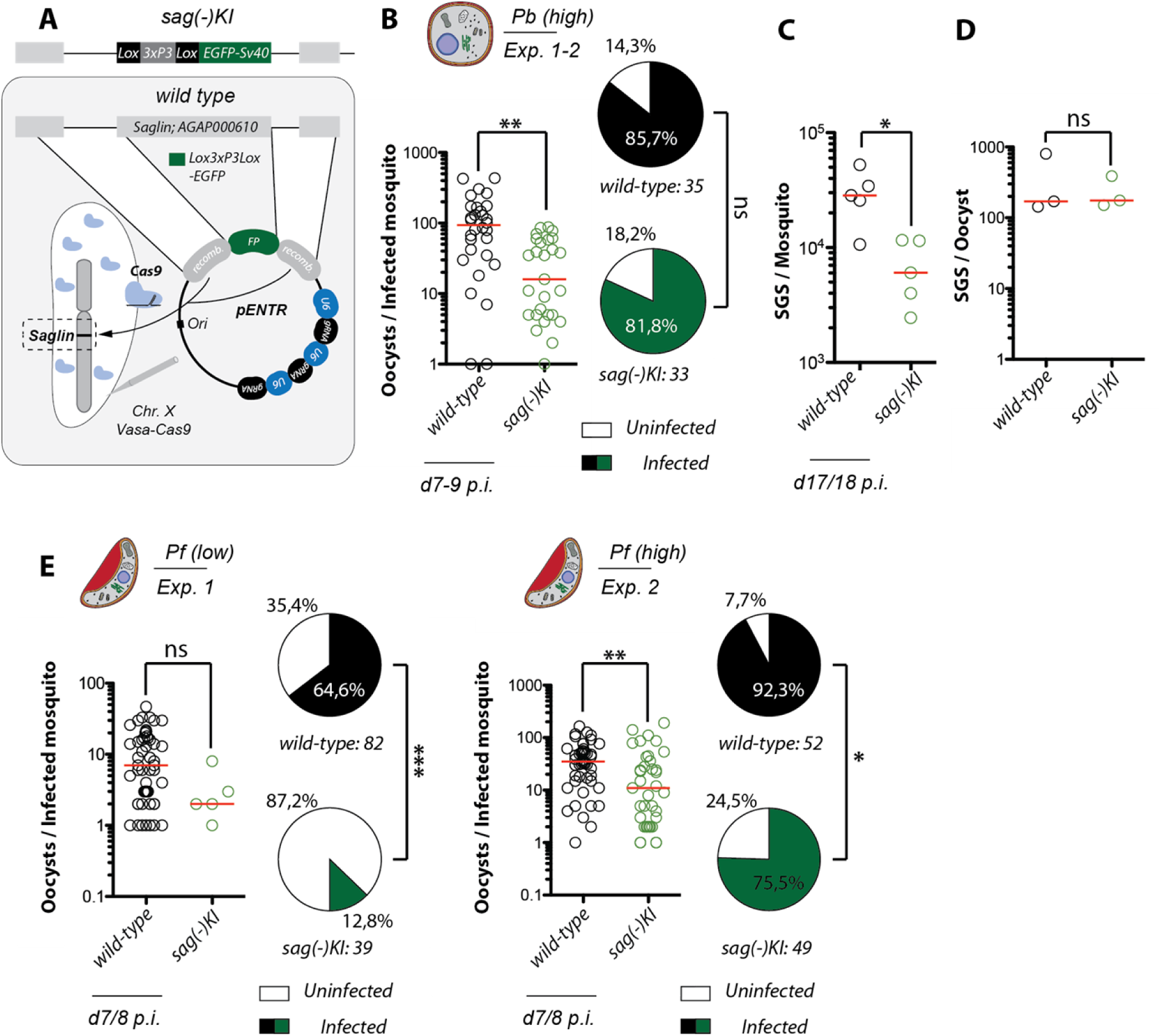
Absence of Saglin impairs midgut colonization by *Plasmodium berghei* and *Plasmodium falciparum*. **A**) Generation of *sag(-)KI* mosquitoes by Cas9 mediated site-directed mutagenesis inducing complete replacement of the *saglin* gene (AGAP000610) with a provided repair template encoding the fluorescence marker EGFP. Please note that the illustration is not drawn to scale. **B)** Oocyst densities in *sag(-)KI* and wild-type siblings infected with the rodent malaria parasite *P. berghei* (*Pb*). Pooled results of two independent experiments. The red line indicates the median. Mann Whitney test: **p = 0.0013. Pie charts represent prevalences of infection. Fisher’s exact test: not significant (ns). **C)** Counts of salivary gland sporozoites (SGS) 17-18 days post infection. Data represent five countings obtained from three infection experiments. Shown is the mean and the standard error of the mean (SEM) of five countings obtained from three different infections. Mann Whitney test: *p = 0.0317. **D)** SGS to oocyst ratio calculated based on paired experiments where both oocyst and salivary gland sporozoites were counted. Mann Whitney test: not significant (ns). **E)** Quantification of oocyst densities and prevalence of infection in *sag(-)KI* and control mosquitoes infected with *P. falciparum* in two different experiments. Data are presented as in B. Mann Whitney test on oocyst densities. Exp. 1: not significant, p = 0.0751; Exp. 2: **p = 0.0034. Fisher’s exact test on prevalence of infection. Exp. 1: ***p < 0.0001 ; Exp. 2: *p = 0.0286

### Absence of Saglin expression has no effect on blood feeding behavior but impairs gut colonization by *Plasmodium*

Functional depletion of saliva proteins can trigger severe effects on the blood feeding behavior in *Anopheles* mosquitoes. For example, co-expression of a single-chain antibody against AAPP reduces the proportion of blood-feeding females as well as hemoglobin uptake, resulting in a lower number of eggs laid per female (Islam *et al*., 2019). Impaired blood uptake could thus be a possible explanation for reduced midgut colonization with *Plasmodia*, as a reduction of the ingested blood volume would likely correlate with a reduction in the number of uptaken gametocytes. To test whether loss of Saglin affects blood feeding, homozygous *sag(-)KI* and wild-type females, bred either as a mixture or separately but always synchronously, were allowed to take blood on the arm of a human volunteer for ten minutes. Subsequently, blood-fed and non-fed females were counted for each genotype. Feeding rates of *sag(-)KI* and WT females were similar and >90% (**Fig. 3A**). We next examined whether blood-fed *sag(-)KI* females ingested the same amount of blood as WT. For this, 155 *sag(-)KI* and 156 WT blood-fed females were weighed using a precision balance. *Sag(-)KI* females were in fact significantly heavier compared to *control* females after blood feeding (**Fig. 3B**). Unlike females of other mosquito species such as *Aedes spp.*, *Anopheles spp*. females raise the hematocrit of their blood meal by excreting fluid during blood ingestion. Hence blood meals of similar volumes may have different concentrations of red blood cells. Thus we also quantified the number of red blood cells in the blood meals of *sag(-)KI* and WT females using a hemocytometer. The cell numbers in blood meals of *sag(-)KI* females is significantly higher compared to WT females (**Fig. 3C**), consistent with the observation that *sag(-)KI* females seem to uptake a greater volume of blood (**Fig. 3B**). To assess possible effects due to the altered blood uptake behavior or potential other consequences of the lack of Saglin on the fitness of *sag(-)KI* mosquitoes, we investigated the persistence of the *sag(-)KI* transgene in competition with WT. This was done by crossing females homozygous for the *sag(-)KI* transgene with WT males. The ratio of individuals positive and negative for the *sag(-)KI* transgene was tracked by flow cytometry over 10 generations, revealing that the frequency of transgene carriers and non-carriers corresponds to the Hardy-Weinberg equilibrium for X-linked inheritance (**Fig. 3D**). Taken together these results suggest no impairment of fitness in *sag(-)KI* mosquitoes, and no impairment of blood feeding behavior in *sag(-)KI* females, ruling out this possible explanation for reduced infection by *Plasmodia*. Therefore we investigated potential differences in the developmental progression of *P. berghei* in the mosquito midgut after blood feeding. The only *Plasmodium* stages able to develop in the mosquito midgut upon blood feeding are female and male gametocytes. Activated male gametocytes divide in 8 microgametes that can easily be visualized using the same staining protocols as for asexual blood stages (**Fig. 3E**). Here we determined the microgametocytemia in individual mosquito blood meals 10-12 minutes after ingestion. Mosquitoes we fed on highly (*Pb high*) and lowly (*Pb low*) parasitized mice. Quantification of blood meals obtained from *Pb low* infected mosquitoes revealed a significant decrease in the microgametocytemia of *sag(-)KI* compared to WT females (**Fig. 3F**). In contrast, no significant difference in the number of microgametes was observed in *Pb high* infected mosquitoes although the same tendency was observed (**Fig. 3F**). To determine whether differences may also be visible at the ookinete stage, the number of transmigrated ookinetes visible on the basal side of the midgut was determined ∼24 hours after blood feeding. In line with the decreased number of microgametes in *sag(-)KI* females, numbers of traversed ookinetes were significantly reduced in both *Pb low* and *Pb high* infected *sag(-)KI* mosquitoes (**Fig. 3G**). Taken together, our results suggest that Saglin plays a role in the development of *Plasmodium* parasites in the mosquito midgut, and we thus made use of the *sag(-)KI* as a reporter line to examine precisely Saglin expression pattern.

**Figure 3.**
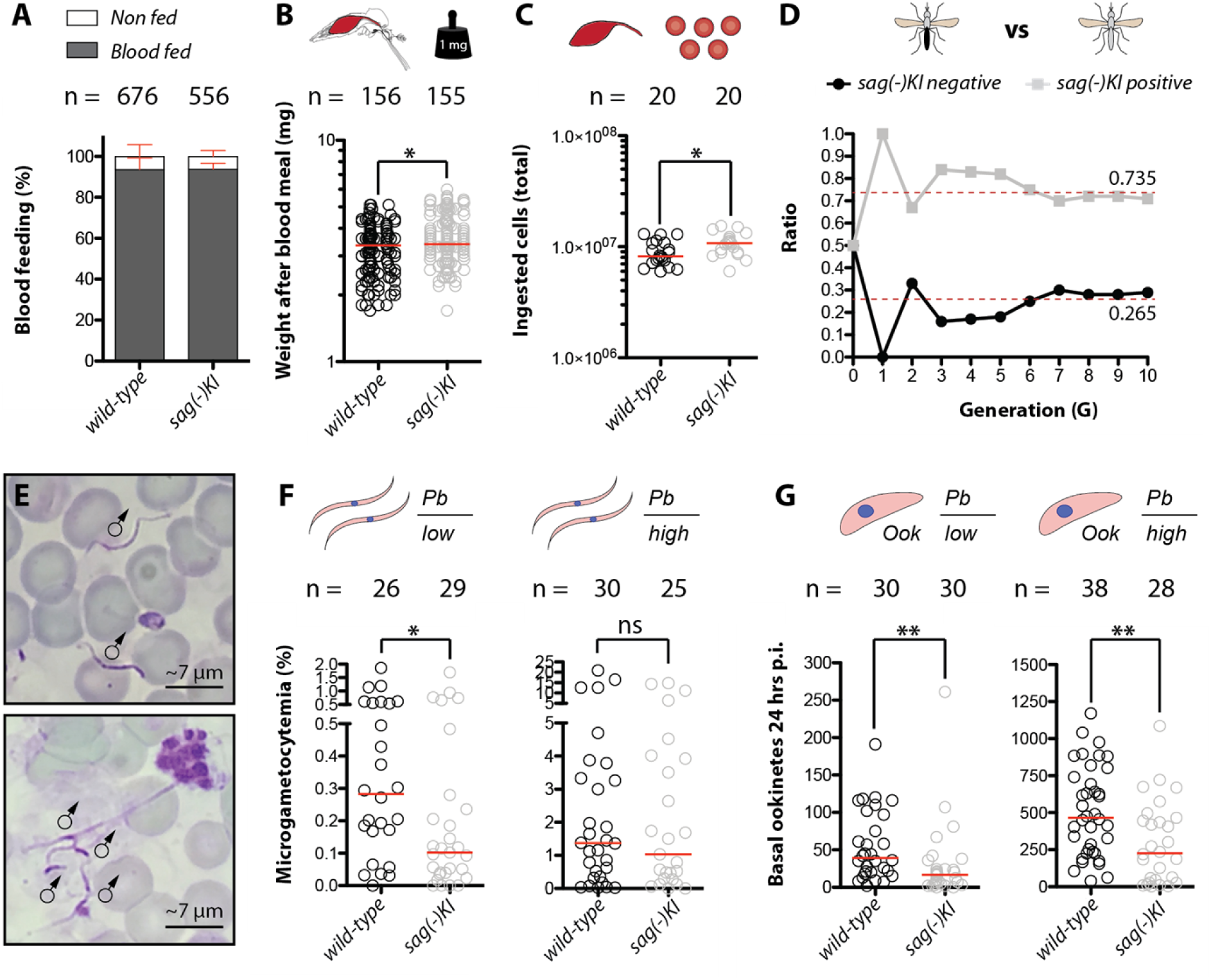
Absence of Saglin impairs exflagellation and reduces the number of ookinetes successfully traversing the midgut but has no detectable fitness costs. **A)** Percentage of blood fed *control* and *sag(-)KI* females after 10 minutes of feeding. Mean and standard deviation (SD) of four independent experiments. **B)** Mosquito weight after blood feeding and red blood cell count (#RBC) in the blood meal (**C**) after 10 minutes of blood feeding of wild-type (*Ngousso)* and *sag(-)KI* females. Data represent pooled results of four experiments. Red line indicates the median. Experiments were performed ≥4 days after eclosure. Mann Whitney test: *p = 0.0133 (weight); *p = 0.0248 (#RBC). **D)** Presence of the *sag(-)KI* transgene in a colony which has been established by crossing homozygous sag(-)KI females to wild type males (F0). The proportion of mutant (homozygote + heterozygous females and hemizygous males) and wild-type was tracked by flow cytometry (COPAS) of neonate larvae at each generation, using the 3xP3-GFP fluorescence marker disrupting the *saglin* coding sequence. Dotted red lines indicate the expected ratios for an X-linked allele in Hardy-Weinberg equilibrium. **E)** Diff-Quik stained smears of mosquito blood meals 10-12 minutes after feeding. Exflagellated male gametes are visible as elongated spirochete-like cells. Scale bar: ∼7 µm. **F)** Numbers of exflagellated male microgametes in the blood meal 10-12 minutes after feeding and number of ookinetes (**G**) visible on the basal side of the midgut epithelium of wild-type (*Ngousso)* and *sag(-)KI* mosquitoes from two independent experiments each. Mosquitoes were infected using lowly (*Pb low*) or highly (*Pb high*) parasitized mice. Red lines indicate the median. Mann Whitney test: *Pb low*: *p = 0.0315; *Pb high*: not significant, p = 0.8327 (# of microgametes). *Pb low*: **p = 0.0014; *Pb high*: **p = 0.0036 (# of basal ookinetes).

### Expression of Saglin is female-specific and restricted to the median lobes of the salivary glands

In *sag(-)KI* mosquitoes, the *EGFP* gene replacing the *saglin* coding sequence (**Fig. 2A**), is placed under the control of the *3xP3* (driving expression in the eyes and nervous system) and *saglin* promoters, and functions as a reporter for transcriptional activity of the *saglin* promoter. The presence of both promoters upstream of *EGFP* thus leads to a *3xP3*-specific expression in the nervous system (in adults, mainly in the eyes) and a *saglin*-specific expression in the salivary glands. We created a second saglin knockout line called *sag(-)EX*, in which the *3xP3* promoter was removed by Cre-mediated excision, thus leaving *EGFP* under the control of the *saglin* promoter exclusively (Klug *et al*., 2021). We could show that the *3xP3* promoter also affects the activity of the endogenous *saglin* promoter, by increasing reporter expression levels in the median lobe (Klug *et al*., 2021). To investigate *saglin* promoter activity in salivary glands and midgut, we thus used the *sag(-)KI* line displaying the same expression pattern as *sag(-)EX* but giving stronger fluorescence signals, and the *sag(-)EX* when necessary to confirm the profile. In *sag(-)KI*, *EGFP* is strongly expressed in the salivary glands, to the point of being visible through the cuticle of the mosquito (**Fig. 4A**). Promoter activity was observed in females but completely absent in males, indicating that Saglin is female-specific (**Fig. 4A**) as previously reported for the mRNA transcript (Arcà *et al*., 2005). Dissections of salivary glands obtained from *sag(-)KI* females revealed that the *saglin* promoter is only active in the median lobe of each salivary gland, with no *EGFP* expression in the proximal- and distal-lateral lobes (**Fig. 4B**). This very specific expression pattern became evident by crossing *sag(-)KI* mosquitoes with the salivary gland reporter lines *trio-DsRed* and *aapp-DsRed* expressing *DsRed* exclusively in the median and distal lobes, respectively (Klug *et al*., 2021). Accordingly, the EGFP signal colocalized with DsRed exclusively in the salivary glands obtained from females heterozygous for *sag(-)KI* and *trio-DsRed*, but not *aapp-DsRed* (**Fig. 4B**). As blood feeding can specifically induce expression of genes important for blood digestion in the midgut (Nolan *et al*., 2011) where Saglin could potentially directly interact with *Plasmodium* parasites ingested by blood feeding, we assessed *EGFP* expression in the midgut 24h after blood feeding using the *sag(-)EX* line (**Fig. 4C**). Indeed, we noticed EGFP fluorescence in cells of the pyloric valve of sugar fed *sag(-)KI* midguts. Other transgenic mosquito lines expressing fluorescent reporters under control of the *3xP3* promoter also presented this fluorescence pattern, suggesting that it is *3xP3*-specific. Indeed, no EGFP signal was detected in blood-fed and sugar-fed *sag(-)EX* midguts, while blood-fed midguts of the *A. stephensi* line *As-G12* expressing GFP under control of the blood meal inducible *G12* promoter showed strong and uniform GFP expression (**Fig. 4C**) (Nolan *et al*., 2011). Taken together, our results indicate that Saglin is specifically expressed in the salivary gland median lobes of female mosquitoes.

**Figure 4.**
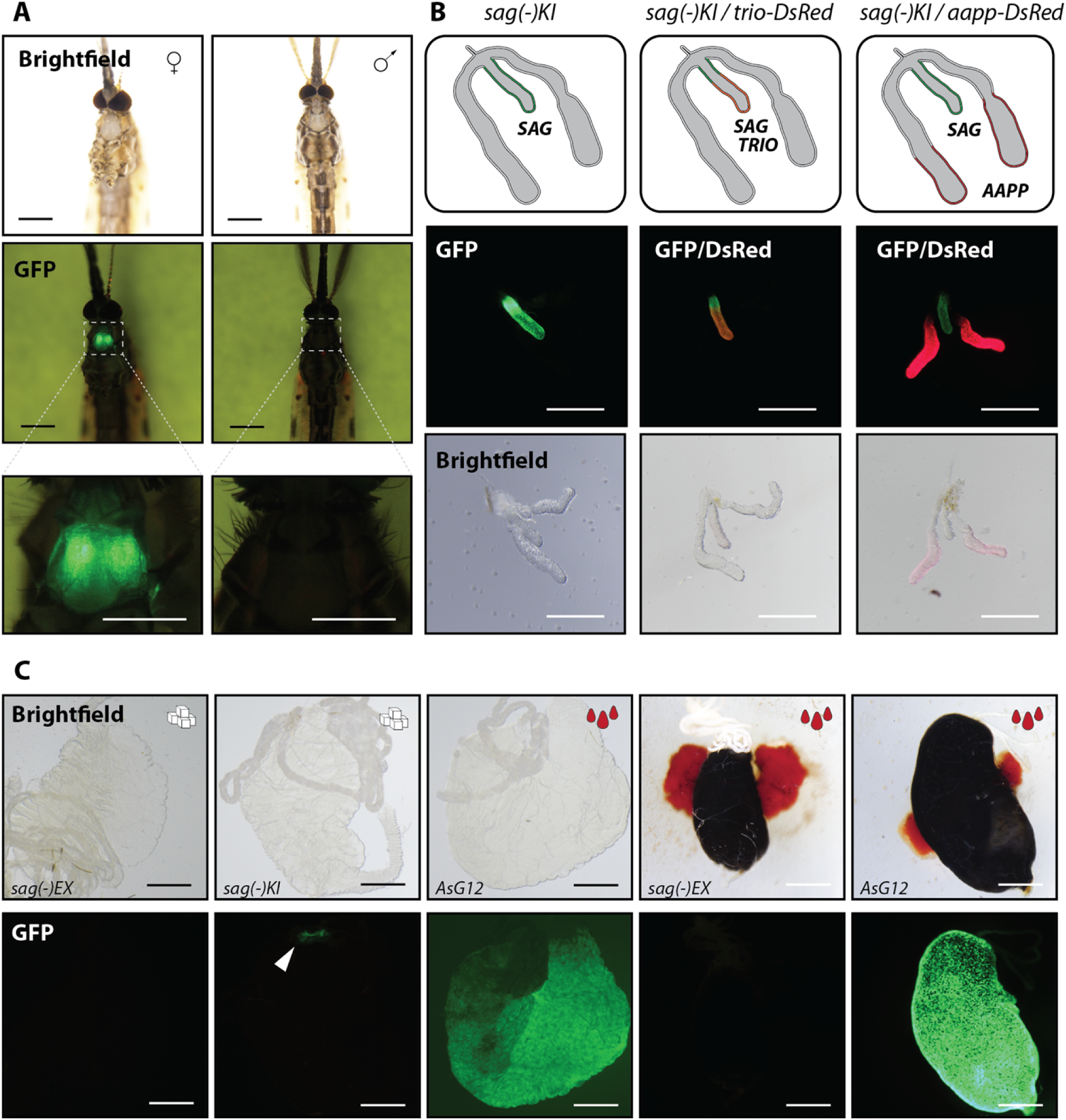
Saglin is exclusively expressed in the median lobes of the salivary glands of female mosquitoes. **A)** Brightfield images (top) and GFP signal in a homozygous *sag(-)KI* female and male mosquito. Scale bar: 0.5 mm. Close-ups of the GFP signal in the thorax are shown below. Scale bar: 250 µm. Male and female mosquitoes were imaged seven days after emergence using the same settings. **B)** *Sag(-)KI* females were crossed to males of the salivary gland reporter lines *trio-DsRed* and *aapp-DsRed*. Salivary glands of heterozygous F1 females were dissected and imaged. The genotypes as well as the expected expression patterns within the salivary glands are shown on the top. Fluorescence (middle) and brightfield (bottom) images of the same salivary glands. Scale bar: 250 µm. **C)** Comparison of EGFP expression in a sugar fed *sag(-)KI* midgut compared to sugar and blood fed *sag(-)EX* midguts. As positive control midguts of the *A. stephensi* line *AsG12* which expresses EGFP under control of the blood meal inducible promoter G12 are shown (Nolan *et al*., 2011). Adults were dissected 7-10 days post emergence. Displayed *AsG12* midguts were dissected either four days after blood feeding (blood already digested) or 24hrs after blood feeding (blood filled). The white arrow indicates the EGFP expressing pyloric valve in *sag(-)KI* which is based on gene expression driven by the 3xP3 promoter. Scale bar: 250 µm.

### Saglin is expressed in the salivary glands, excreted during blood feeding and re-ingested with the blood meal

A previous study presented evidence that Saglin might function as a receptor mediating recognition and invasion of the salivary glands by *Plasmodium* sporozoites (Ghosh *et al*., 2009). To investigate whether Saglin is exposed on the salivary gland surface to come in contact with sporozoites and whether Saglin can spread to the distal lobes of the salivary gland (distal lobes are more infected by sporozoites than the median lobe), we performed immunofluorescence experiments using α-Saglin antibodies. Similar to the *EGFP* expression in *sag(-)KI* mosquitoes, saglin-specific staining of control salivary glands was observed in the acinar cells of the median lobe (**Fig. 5A**). Of note, due to the strong polarity of the acinar cells that form an inward cavity towards the salivary duct, their cytoplasm is restricted to the edges of the median lobe. Accordingly, saglin-specific signals were observed in the marginal area, likely present in the acinar cell cytoplasm although we cannot exclude the possibility that a subset of Saglin is localised on the surface of the median lobe. In addition, we observed very strong Saglin signals in the lumen of the proximal region of lateral lobes (**Fig. 5A**), although cells in these regions did not show any *saglin* promoter activity (**Fig. 4B**). This observation is in line with previous findings showing a similar immunofluorescence staining pattern for Saglin in *A. stephensi* (O’Brochta *et al*., 2019). Interestingly, fluorescence signals in the proximal lobes were significantly stronger than in the median lobe, which could indicate differences in protein concentrations, or in the antibody-mediated recognition of Saglin when the protein is intracellular or secreted. As expected, salivary glands obtained from homozygous *sag(-)KI* females displayed no Saglin signal (**Fig. 5A**). *Plasmodium* sporozoites have been reported to preferentially invade the distal lobes (Wells and Andrew, 2019). To investigate a possible interplay between Saglin expression in the median lobe and physical accessibility through the hemolymph that would allow sporozoites to sense and make contact with the salivary gland, we have injected XFD594 labeled wheat germ agglutinin (WGA) into live mosquitoes. Imaging of 13 dissected salivary glands revealed a uniform staining of distal and proximal lobes while >50% of the median lobes were devoid of WGA coating (**Fig. 5B, Fig. 5 – figure supplement 1**). When median lobes displayed fluorescence, the signal was either weak compared to distal lobes, or restricted to the tip (**Fig. 5 – figure supplement 1**). Of note, and in contrast to lateral lobes, the median lobe was often tightly imbedded in fat body tissues, which may in part explain its lower accessibility. WGA staining was specific upon injection and no difference in the staining pattern was observed between wild-type and *sag(-)KI* mosquitoes (**Fig. 5 – figure supplement 2**). To further confirm the salivary gland-specific activity of the *saglin* promoter and fate of Saglin, we performed a series of western blot experiments on dissected tissues (**Fig. 5C**). Staining with α-Saglin antibodies revealed a strong band within samples of median lobes and faint bands in samples of lateral lobes obtained from WT. No Saglin was detectable in salivary gland samples obtained from *sag(-)KI* mosquitoes (**Fig. 5C**), confirming that the mutant is a full knockout. Similarly, EGFP was highly enriched in the median lobe sample of *sag(-)KI* mosquitoes with a fainter signal in lateral lobes (**Fig. 5C**). In contrast, no Saglin was detected in midguts (**Fig. 5C**), hemolymph and carcasses deprived of salivary glands of sugar fed mosquito females (**Fig. 5D**), likely excluding any *saglin* expression in other tissues as well as Saglin dissemination originating from the salivary glands. Saglin was detected in proboscis samples collected from WT but not *sag(-)KI* females (**Fig. 5E**), suggesting that Saglin is a component of saliva and potentially excreted during salivation. In line with these observations, a previous study using mass spectrometry has detected Saglin in the saliva of mosquitoes (Okulate *et al*., 2007). Thus we hypothesized that Saglin might be ingested during blood sucking. Indeed, we could detect Saglin in the stomach contents of blood-fed but not sugar-fed WT females, revealing that Saglin can indeed be present in the gut, but only after blood feeding (**Fig. 5F**). Taken together, our data thus suggest that Saglin is specifically expressed in the median lobe of the salivary gland, probably stored in the duct of the proximal-lateral lobes and injected in the host skin upon salivation, where it can be ingested by the mosquito during blood sucking.

**Figure 5.**
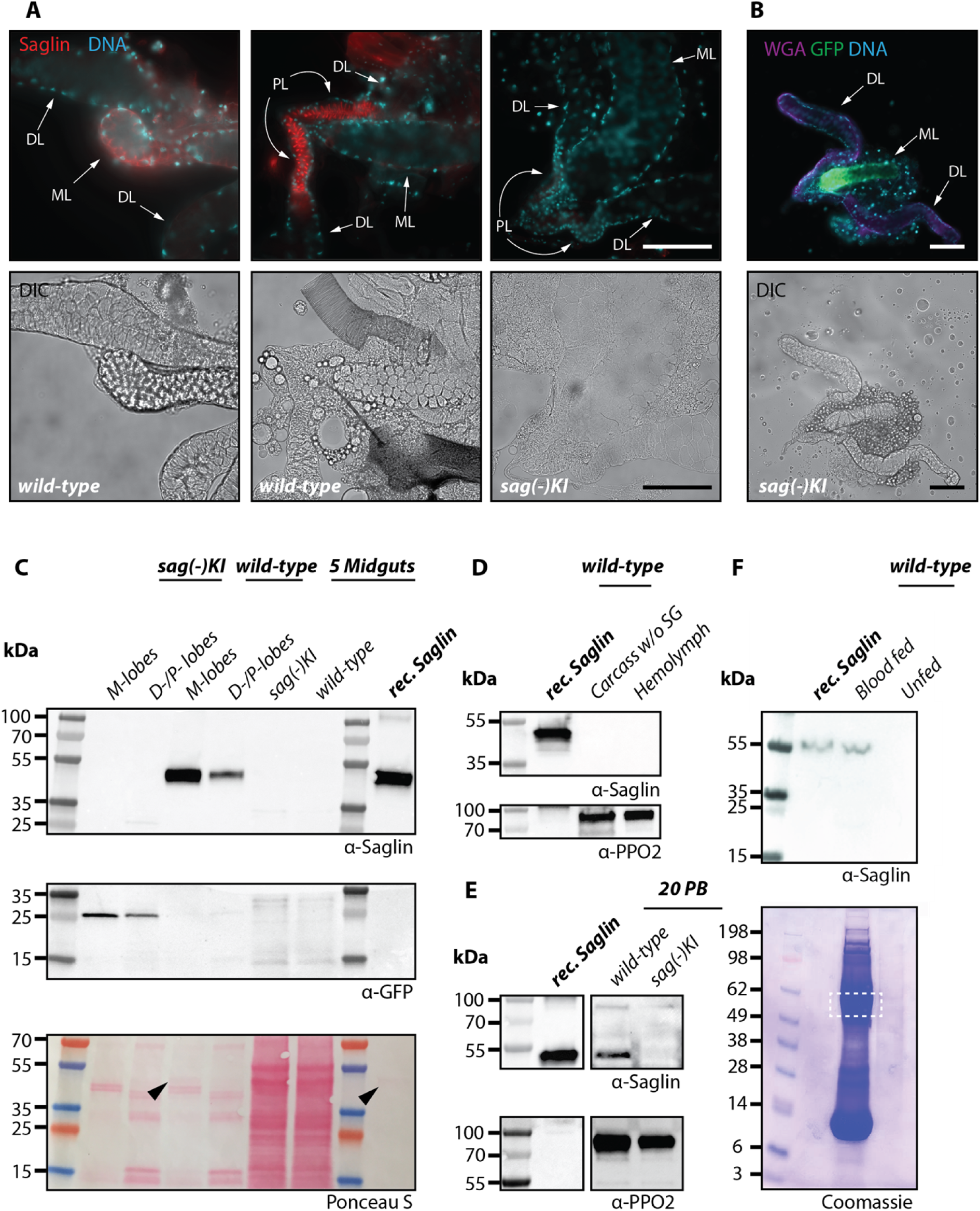
Saglin is a protein found in the salivary glands that is excreted with saliva and reabsorbed with the blood meal. **A)** Immunofluorescence staining of Saglin in fixed wild-type (*Ngousso*) and *sag(-)KI* salivary glands. Saglin (red) and DNA (blue) staininges (top), and differential interference contrast (DIC, bottom). Images depict the same salivary glands. DL: distal-lateral lobes; PL: distal-proximal lobes; M: median lobe. Scale bar: 100 µm. **B)** Staining with fluorophore coupled WGA of a *sag(-)KI* salivary gland *ex vivo*. WGA staining in purple, DNA in cyan and *saglin* promoter based expression of EGFP in green. Scale bar: 100 µm. **C)** Western blots of different mosquito tissues to detect the presence of Saglin. Median (M-lobes) and Distal-lateral/distal-proximal lobes (D/P-lobes) in comparison to unfed midguts of the same wild-type (*Ngousso*) and *sag(-)KI* mosquitoes, carcasses with salivary glands removed and hemolymph of wild-type (**D**), proboscis of wild-type and *sag(-)KI* (PB) (**E**) and stomach contents of sugar and blood fed wild-type midguts. Note that the blot shown in E is shown as two different images because of different exposure bands and removed lanes in between. (**F**). Recombinant Saglin was used as positive control and western blots and SDS-Page gels were treated with α-PPO2 antibodies, Ponceau S or Coomassie blue, respectively, to ensure equal loading of samples. Arrows in C indicate Ponceau S stained Saglin present in median lobes of wild-type and the recombinant protein control. Dashed square in F indicates the area Saglin would be expected to localize in the Coomassie stain.

### Priming of the skin bite site with wild-type mosquitoes does not rescue *Plasmodium* infection in *sag(-)KI* mosquitoes

Therefore, we hypothesised that Saglin ingested with the blood meal may have a pro-parasitic effect on *Plasmodium* midgut colonization. If so, the observed *Plasmodium* infection reduction in *sag(-)KI* mosquitoes might be rescued by the presence of saliva from wild-type mosquitoes. To test this hypothesis we performed co-feeding and priming experiments by either feeding homozygous *sag(-)KI* and wild-type mosquitoes as a mixture or in a sequential manner on the same *P. berghei* infected mouse (**Fig 6A**). We speculated that when *sag(-)KI* mosquitoes uptake blood at sites where wild-type mosquitoes have bitten before, remaining saliva in the skin may rescue *Plasmodium* infection in *sag(-)KI* mosquitoes. We performed three to four independent experiments and normalized results to the mean of the oocyst loads in WT mosquitoes for each experiment (**Fig 6B**). Oocyst ratios and infection prevalences of *sag(-)KI* mosquitoes remained inferior to those of WT controls but the difference was not affected by the feeding regime tested (**Fig. 6B,C**), suggesting that wild-type priming cannot rescue Saglin deficiency during *Plasmodium* infection, possibly because the protein is not stable or available long enough at the bite site.

**Figure 6.**
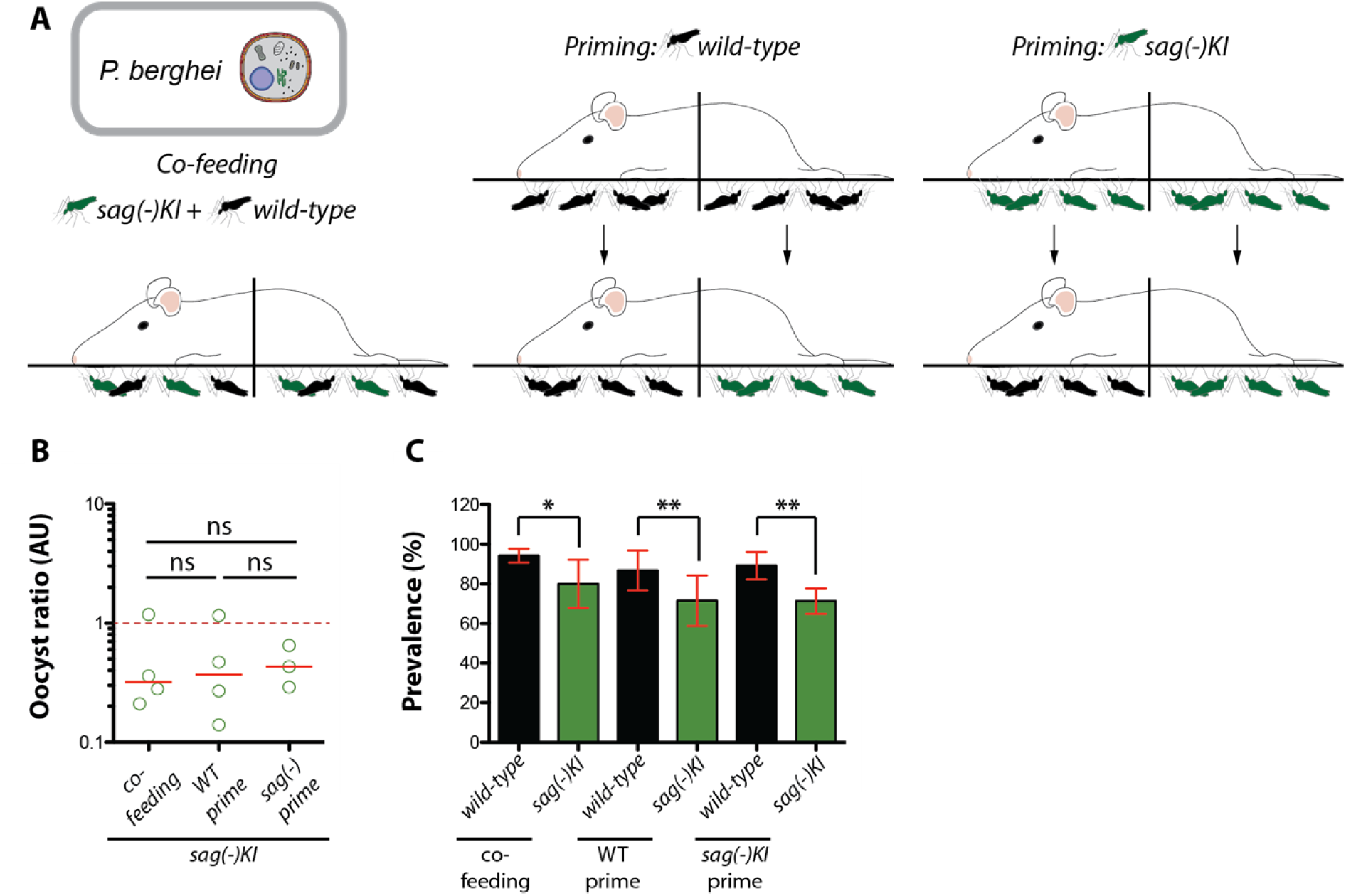
Priming of the bite site has no effect on the infection phenotype in *sag(-)KI* mosquitoes. **A)** Co-feeding of *sag(-)KI* and wild-type mosquitoes on the same *P. berghei* infected mouse, and infections of *sag(-)KI* and wild-type mosquitoes on mice primed with wild-type and *sag(-)KI*. **B)** Oocyst ratios of *sag(-)KI* mosquitoes which have been infected using conditions as illustrated in A. Data represent 3-4 independent experiments. The solid red line indicates the median, while the dashed red line indicates the value at which *sag(-)KI* and wild-type mosquitoes reach equilibrium in parasite load. one-way-ANOVA (Kruskal-Wallis test): ns: not significant. **C)** Prevalence of infection for experiments shown in B. Fisher’s exact test: *p = 0.0293 (co-feeding); **p = 0.002 (WT prime); **p = 0.0031 (*sag(-)KI* prime).

### Saglin is essential for successful transmission at low infection densities

To test whether the decrease in parasite load observed *in sag(-)KI* mosquitoes could affect mosquito capability to transmit *Plasmodium* sporozoites, we performed mouse infection experiments using wild-type and *sag(-)KI* mosquitoes infected with *P. berghei*. Because *P. berghei* infections in *Anopheles spp.* mosquitoes generally cause a much higher parasite load compared to *P. falciparum*, two different infection regimes were chosen to achieve high and low parasite loads while maintaining high prevalence. Highly infected mosquitoes (*Pb high*) reflect common *P. berghei* infection levels achieved under laboratory conditions, while mosquitoes with a low parasite load (*Pb low*) reflect parasite numbers more commonly seen in mosquitoes infected with *P. falciparum* in the wild. For this, mosquitoes were infected from mice showing a parasitemia of either <1% (*Pb low*) or >2% (*Pb high*). *Pb high*-infected wild-type and *sag(-)KI* mosquitoes displayed a transmission efficiency of 100%. Of the 6 mice bitten by *Pb high*-infected wild-type mosquitoes, all became positive after a prepatent period of 4.7 days. Similarly, 4 of 4 mice bitten by *Pb high*-infected *sag(-)KI* mosquitoes became positive after a prepatent period of 4 days (**Fig. 7A,C**). Consistent with the observed difference of 0.7 days in prepatency, mice infected by *sag(-)KI* mosquitoes showed faster parasite growth and, on average, higher parasitemia at day 6 post infection compared with mice infected by wild-type mosquitoes (**Fig. 7A,C**). In contrast, while 6 of 6 mice bitten by *Pb low*-infected wild-type mosquitoes became positive for blood stage parasites after a prepatent period of 5.3 days, only 1 of 4 mice bitten by *Pb low*-infected *sag(-)KI* mosquitoes became parasite positive after 11 days indicating a severe impairment in the transmission capacity (**Fig. 7A,C**).

**Figure 7.**
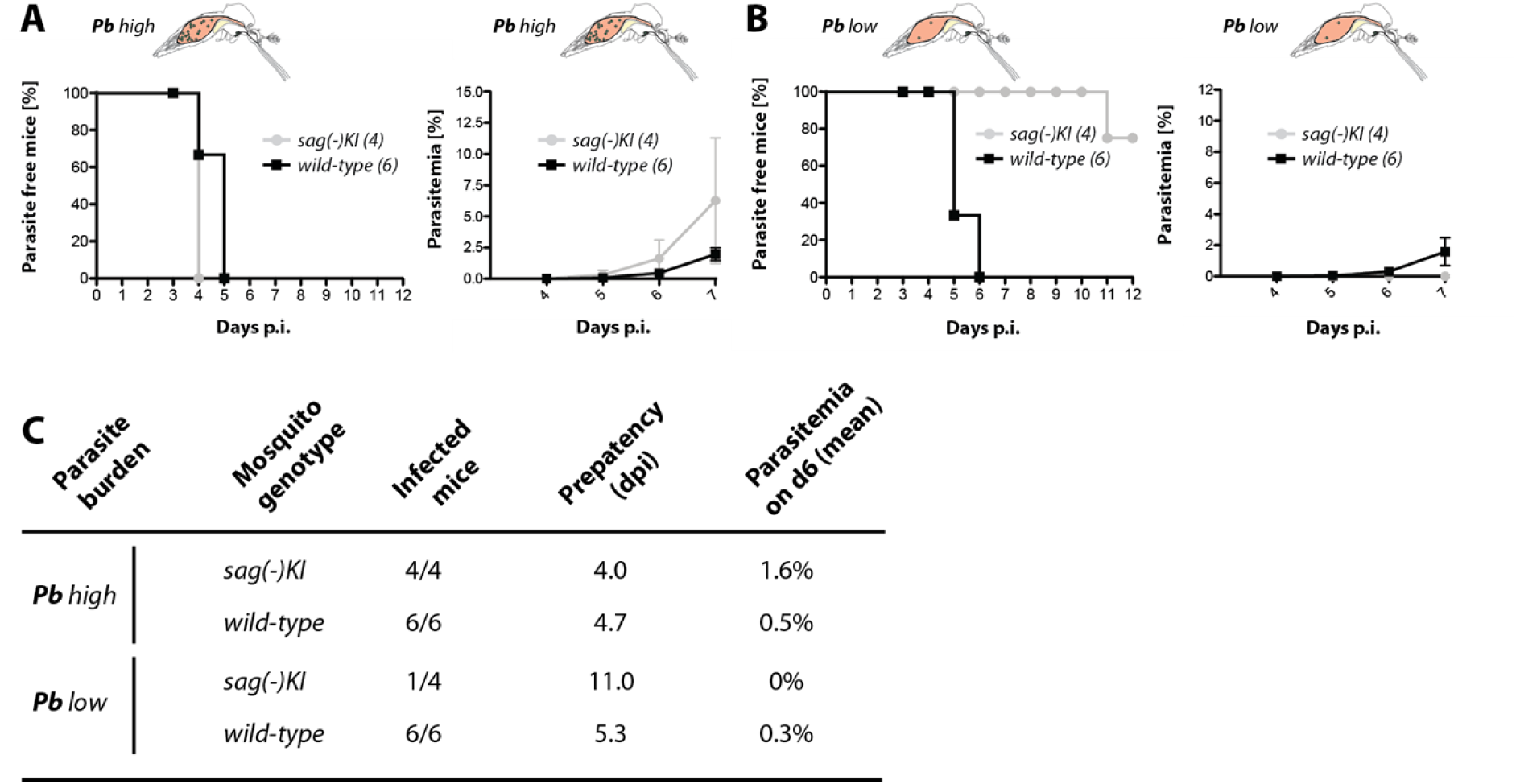
Saglin is required for successfull transmission at low infection densities. **A)** Percentage of parasite free mice after being exposed to the bites of ten *sag(-)KI* or WT mosquitoes with high (*Pb* high) or **B)** low (*Pb* low) parasitemia. The number of mice is given in brackets. The growth of asexual blood stages in infected mice was monitored between day 4 and day 7 after mosquito exposure. **C)** Table summarizing results illustrated in A and B. Given is the mosquito genotype, the numbers for infected / exposed mice as well as the mean prepatency in days and the mean parasitemia at day 6 post infection. Prepatency is defined as the time between mosquito exposure and the first observation of blood stages.

## Discussion

The knockdown of *saglin* has been shown to lower sporozoite burdens in the salivary glands of *P. falciparum* infected mosquitoes, which has been attributed to an impaired interaction with the sporozoite-specific protein TRAP (Ghosh *et al*., 2009). Here we confirm that the complete loss of Saglin in transgenic mosquitoes decreases sporozoite loads in the salivary glands. Still, our data rather support a role of Saglin during parasite development in the midgut, with limited to no contribution to salivary gland invasion by sporozoites. We provide several lines of evidence indicating that Saglin facilitates parasite invasion of the mosquito midgut. Indeed, *sag(-)KI* mosquitoes display a reduced oocyst burden compared to WTs, which may be in part explained by a reduction in male gametocyte exflagellation and in the number of ookinetes able to cross the midgut epithelium. In contrast, we could not detect a specific effect of Saglin knockout on salivary gland invasion by sporozoites. Indeed, the decreased sporozoite burden in *sag(-)KI* mosquitoes was proportional to the decreased oocyst burden. Interestingly, a similar phenotype was recently described for the knockout of the salivary protein SGS1 in *A. aegypti*. Knockout mosquitoes lacking SGS1 expression showed a 48-79% decrease in oocyst numbers upon infection with *P. gallinaceum* (Kojin *et al*., 2021). Immunofluorescence stainings provided here as well as in a previous study performed in *A. stephensi* (O’Brochta *et al*., 2019) demonstrate that Saglin does not localize towards the salivary gland distal lobes where the majority of sporozoites invade (Wells and Andrew, 2019). Saglin could be detected at the edges of the median lobe by immunofluorescence, raising the possibility that a subset of Saglin is exposed at the surface. However, injection of fluorophore-labelled WGA in living mosquitoes stained mainly the distal lobes of the salivary gland, indicating that the median lobe has little contact with hemolymph. In addition, the median lobes are usually surrounded by fat tissue that is difficult to remove during dissection. Taken together, these observations could, at least in part, explain why the distal lobes are preferentially infested by sporozoites (Wells and Andrew, 2019). If this is true, the limited physical access to the median lobe would further limit the ability of surface-exposed Saglin to function as a sporozoite receptor. Furthermore, Saglin is not secreted at detectable levels in the hemolymph where it could potentially serve as a chemotactic attractant for sporozoites. In agreement with the hypothesis that Saglin may not contribute to salivary gland invasion by sporozoites. A recent study has shown that sporozoites do invade the salivary glands of male *Anopheles* mosquitoes (Haraguchi *et al*., 2022) while *saglin* is female-specific (Arcà *et al*., 2005; Klug *et al*., 2021). Taken together, these observations indicate that Saglin is unlikely to be a determinant for sporozoite invasion as sporozoites should not come in contact with this protein before reaching the luminal side of salivary glands. There, TRAP may have a capacity to interact with Saglin. Indeed, TRAP has been shown to bind ubiquitously to a wide range of different proteins such as fetuin-A on hepatocytes (Jethwaney *et al*., 2005), alpha-v-containing integrins (Dundas *et al*., 2018) and platelet derived growth factor receptor β (Steel *et al*., 2021).

Comparison of the expression pattern of *EGFP* in the *sag(-)KI* line with immunofluorescence images revealed that Saglin accumulates at high concentrations in the lumen of the proximal region of the lateral lobes while the *saglin* promoter activity is restricted to the median lobe. This and the fact that Saglin has a signal peptide, suggest that Saglin is secreted into the salivary duct, where it spreads to the proximal lobes, likely to be stored and mixed with other salivary components in preparation for salivation. The notion that Saglin is a component of saliva is supported by mass spectrometric detection of Saglin in saliva samples (Okulate *et al*., 2007). We did not detect Saglin expression (*EGFP* expression in *sag(-)KI* and *sag(-)EX* mosquitoes) or protein (by Western blotting) in all other tissues of female mosquitoes we tested, i.e. hemolymph, midgut, carcasses deprived of salivary glands, including after blood feeding, demonstrating that Saglin is specifically expressed in the salivary glands and resides there. Hence the question, how can Saglin affect parasite development in the midgut if it is solely expressed and resident in the salivary glands? The presence of Saglin in the proboscis and in the gut of WT mosquitoes that had just uptaken blood suggests that Saglin is indeed injected to the host skin at the bite site together with other saliva compounds, and re-ingested together with blood upon sucking. We thus hypothesized that Saglin may interact with parasites at the bite site or in the ingested blood meal. For example, it has been demonstrated for *A. aegypti* that mosquitoes regurgitate much of their own salivary excretions (Whiten *et al*., 2018). Still, priming experiments to allow *sag(-)KI* mosquitoes to uptake traces of Saglin left in the skin by wild-type females, did not rescue *Plasmodium* development to WT levels, possibly because Saglin is not stable long enough or available in high enough quantities to complement Saglin deficiency in *sag(-)KI* mosquitoes.

Should Saglin be important for parasite development in the mosquito midgut, two different possible modes of action are conceivable. Saglin could either function as a cue for gametocytes to become activated and trigger life cycle progression as has been shown for the insect-specific molecule xanthurenic acid (Billker *et al*., 1998). In this scenario, the intrinsic function of Saglin would be uncoupled from its effect on the parasite. The second possibility would be that Saglin alters physiological conditions in the blood meal, which could stimulate *Plasmodium* development. In this case the proparasitic effect of Saglin may not require a direct interaction of the parasite with the protein. No function has yet been assigned to Saglin or any other protein in the SG1 family. Still, we can already draw some conclusions from our data. The knockout of Saglin has no negative effect on blood feeding behavior or on the fitness of mosquitoes under laboratory conditions. In contrast, the suppression of salivary proteins involved in the inhibition of blood coagulation and platelet aggregation, such as the anopheline antiplatelet protein (AAPP) and Aegyptin, have been shown to impair blood feeding behavior and fecundity (Chagas *et al*., 2014; Islam *et al*., 2019). Interestingly, we observed that the infection impairment of *saglin*-knockout mosquitoes applies to both the rodent malaria parasite *P. berghei* and the human malaria parasite *P. falciparum*. While mosquitoes were allowed to feed directly on mice for *P. berghei* infections, *P. falciparum* infections were performed using gametocyte cultures and artificial feeding devices. Of note *P. falciparum* cultures lack cellular components of the immune system and contain only inactivated complement factors. Therefore, Saglin may not be involved in the neutralization of immune factors, which could otherwise have had a protective effect on parasites. No structure of a member of this protein family has been solved, however *in silico* structural prediction of monomeric Saglin using I-Tasser (Roy, Kucukural and Zhang, 2010) revealed that the nucleotide binding domain 1 linked to the middle domain of monomeric Hsp104 from *S. cerevisiae* (PDB: 6AHF) is the closest related structure followed by other chaperones like *Ct*Hsp104 (PDB: 5D4W) and *Tt*ClpB (PDB: 1QVR). Interestingly, the salivary D7 protein from *Culex quinquefasciatus* was shown to bind the nucleotides adenosine diphosphate (ADP) and adenosine triphosphate (ATP) with high affinity. Since ADP and ATP play an important role in activating platelet aggregation, the depletion of both nucleotides in the blood meal by D7 binding enhances blood feeding on mammals (Martin-Martin *et al*., 2020). Other D7 proteins sequester hemostasis mediators such as serotonin and epinephrine with different affinities based on very different ligand specificities (Calvo *et al*., 2009). SG1 proteins share limited sequence similarity with each other (Arcà *et al*., 2005) (**Fig. 8**) but contain several highly conserved residues like C18 and C60 (positions referring to Saglin, AGAP000610) and amino acid stretches (**Fig. 8**). In addition, SG1 proteins are all of similar size (385-431 aa), with Saglin being the largest family member (**Fig. 8-figure supplement 1**). Conserved cysteines together with a similar size might indicate that secondary and tertiary structural elements are at least partially conserved. This could indicate a conserved mode of action of the SG1 family, possibly similar to the D7 proteins that sequester very different molecules, although both protein families do not show significant sequence similarity.

**Figure 8.**
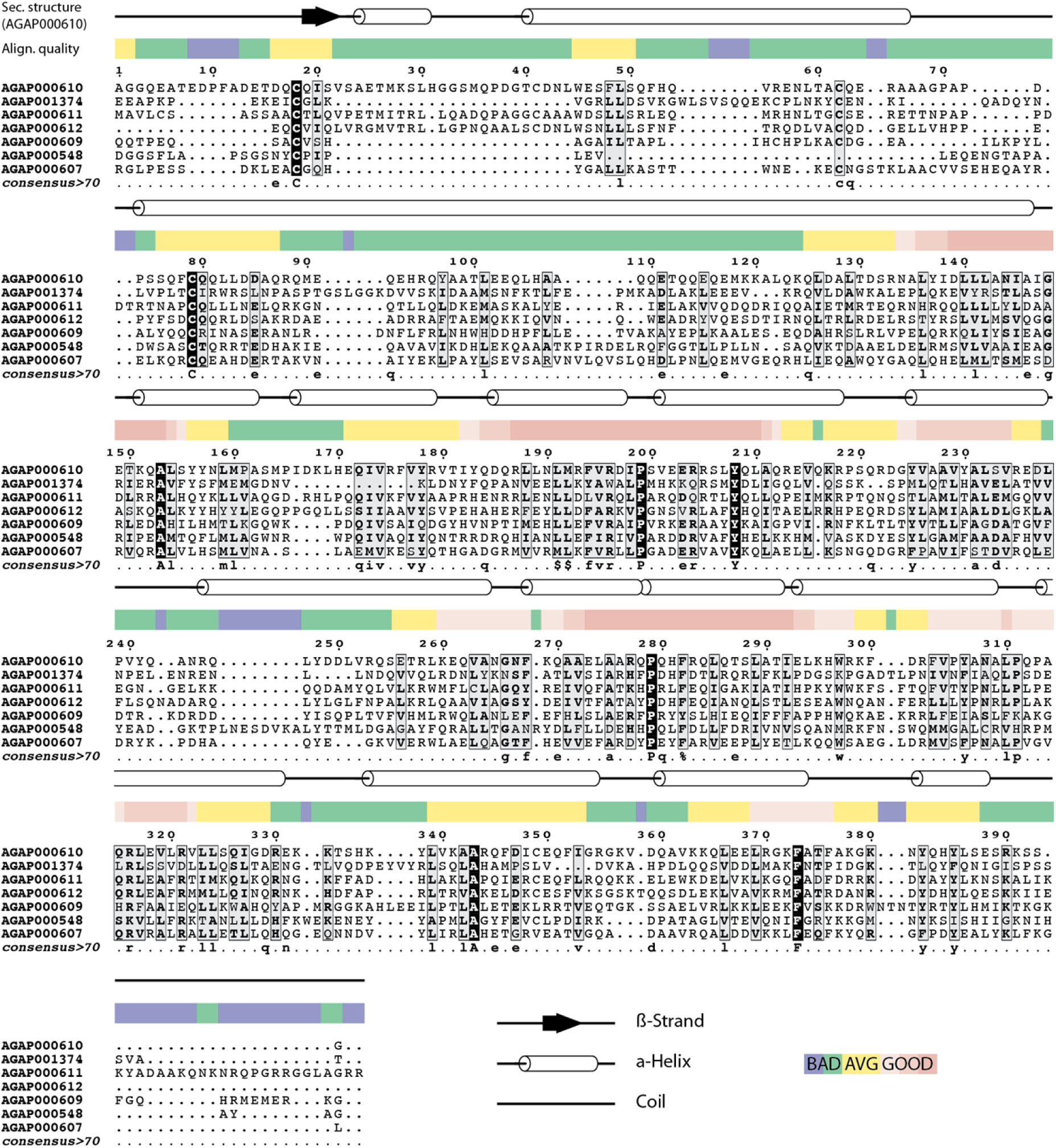
Alignment of SG1 family proteins. Multiple sequence alignment of all seven SG1 proteins known in *Anopheles gambiae.* The initial alignment was performed with PSI-Coffee (Notredame, Higgins and Heringa, 2000) while sequence similarities based on physico-chemical properties of amino acids were calculated with ESPript (Robert and Gouet, 2014). The colored stripe above the alignment indicates the quality of the alignment according to ESPrit. In addition the secondary structure of Saglin (AGAP000610) predicted by I-Tasser is shown along the alignment (Roy, Kucukural and Zhang, 2010). Please note that the signal peptides from all proteins except AGAP000611, for which no signal peptide has been predicted, were removed in preparation of the alignment.

The absence of a fitness cost in Saglin loss-of-function mutants in combination with the strong decrease in transmission capacity for *P. falciparum* (70% decreased oocyst burden) makes Saglin an attractive target for gene drive approaches. Disruption of the *saglin* gene with a driving cassette is expected to decrease the malaria transmission ability of mosquitoes without the need for additional anti-plasmodial effectors. Still, *P. berghei* parasites that managed to invade *sag(-)KI* salivary glands were infectious to mice, thus an additional effector could be introduced to further reduce the transmission capacity of *sag(-)* mosquitoes. We do not know how Saglin affects parasite development in the mosquito midgut, but we showed that Saglin absence reduces the number of microgametes and ookinetes invading the midgut epithelium. Further characterization of Saglin and other SG1 proteins will be instrumental to refine transmission blocking strategies. Should it be possible to uncouple the intrinsic function of Saglin from its effect on the parasite, this could potentially enable the construction of transmission incompetent Saglin alleles retaining their intrinsic function, thereby eliminating the need for gene knockout and reducing the impact of genetic interventions.

---

**Figure 2 – figure supplement 1.**
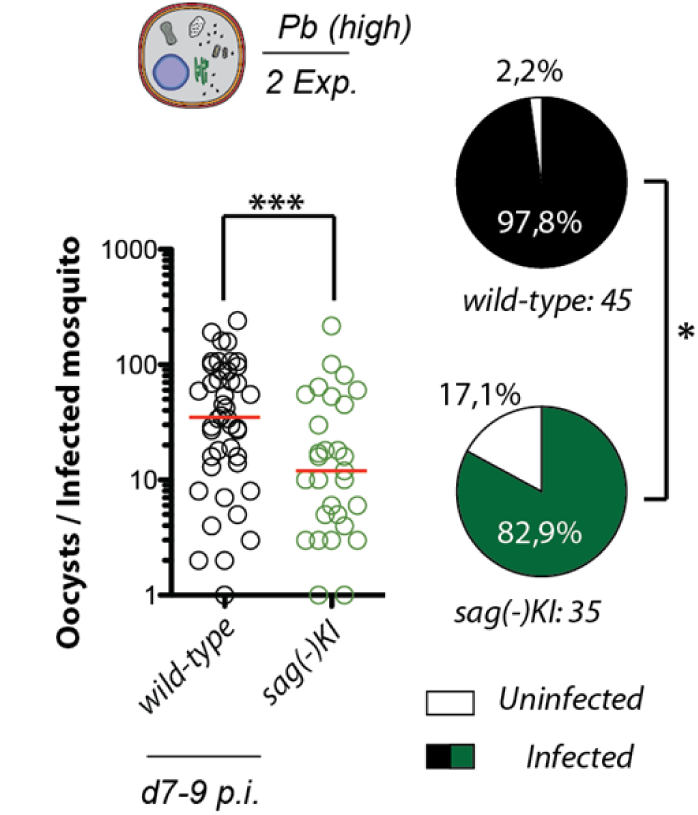
The *sag(-)KI* phenotype is retained even if knockout and wild-type colonies are maintained separately. Oocyst densities in *sag(-)KI* and wild-type (*Ngousso*) mosquitoes derived from two different colonies. Results of two pooled experiments. Mann Whitney test: ***p = 0.0007. Pie charts represent prevalence of infection. Fisher’s exact test: *p = 0.0394.

**Figure 5 – figure supplement 1.**
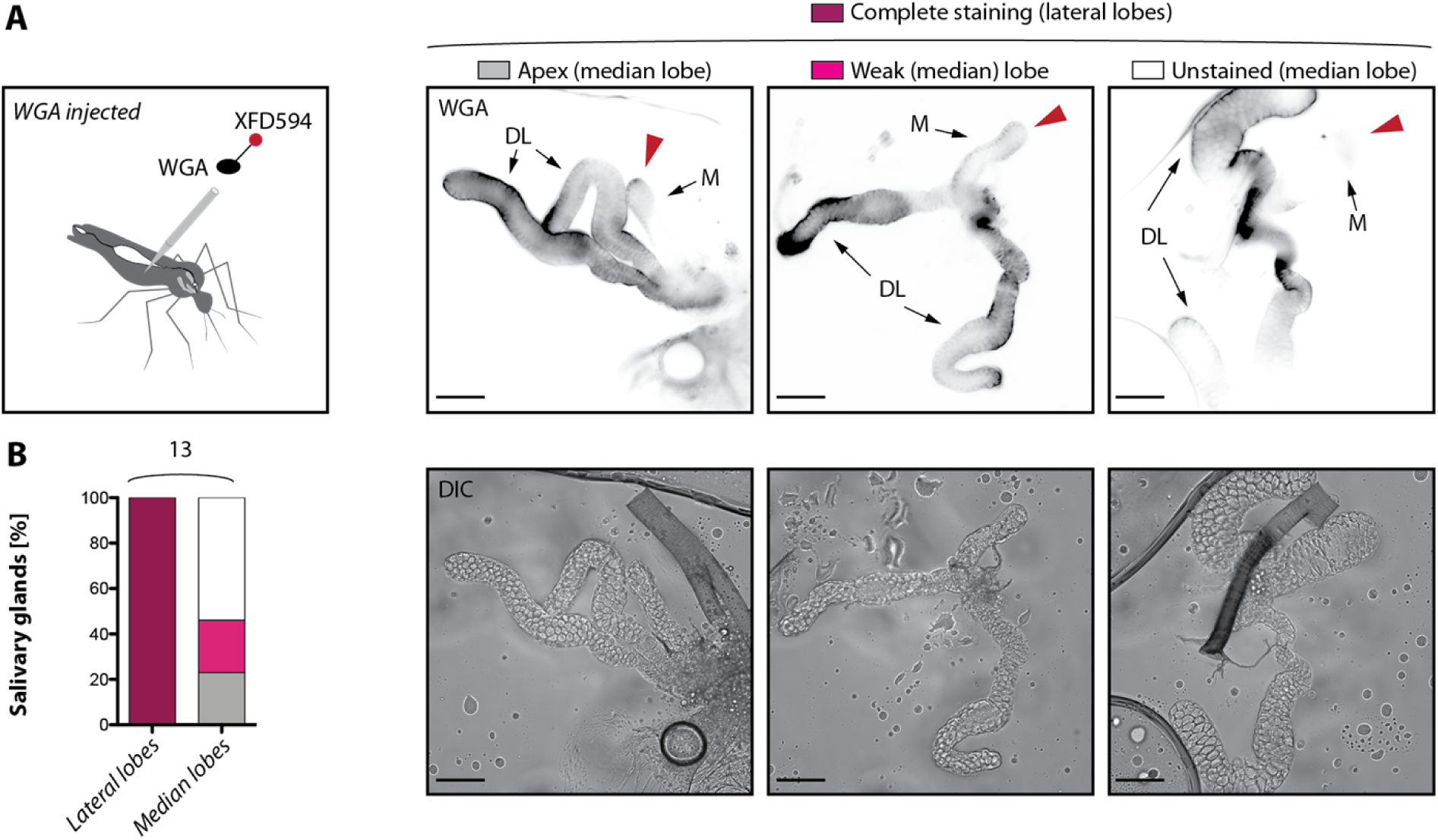
Quantification of *in vivo* WGA staining patterns of median and lateral lobes. **A)** Staining pattern of SGS in mosquitoes injected with wheat germ agglutinin (WGA) coupled to XFD594. WGA-XFDS594 was injected into living mosquitoes using a capillary and mosquitoes were dissected one hour after injection. The observed staining pattern were classified into „complete staining“ for the distal lobes (DL), which was the only pattern observed for this part of the gland, and „apex“, „weak“, „unstained” for the median lobe (M). Images illustrating each pattern are given showing localisation of WGA-XFD594 (top) and differential interference contrast (DIC, bottom). Scale bar: 100 µm. **B)** Quantification of staining patterns according to (A). The number of analyzed salivary glands is indicated above the graph.

**Figure 5 – figure supplement 2.**
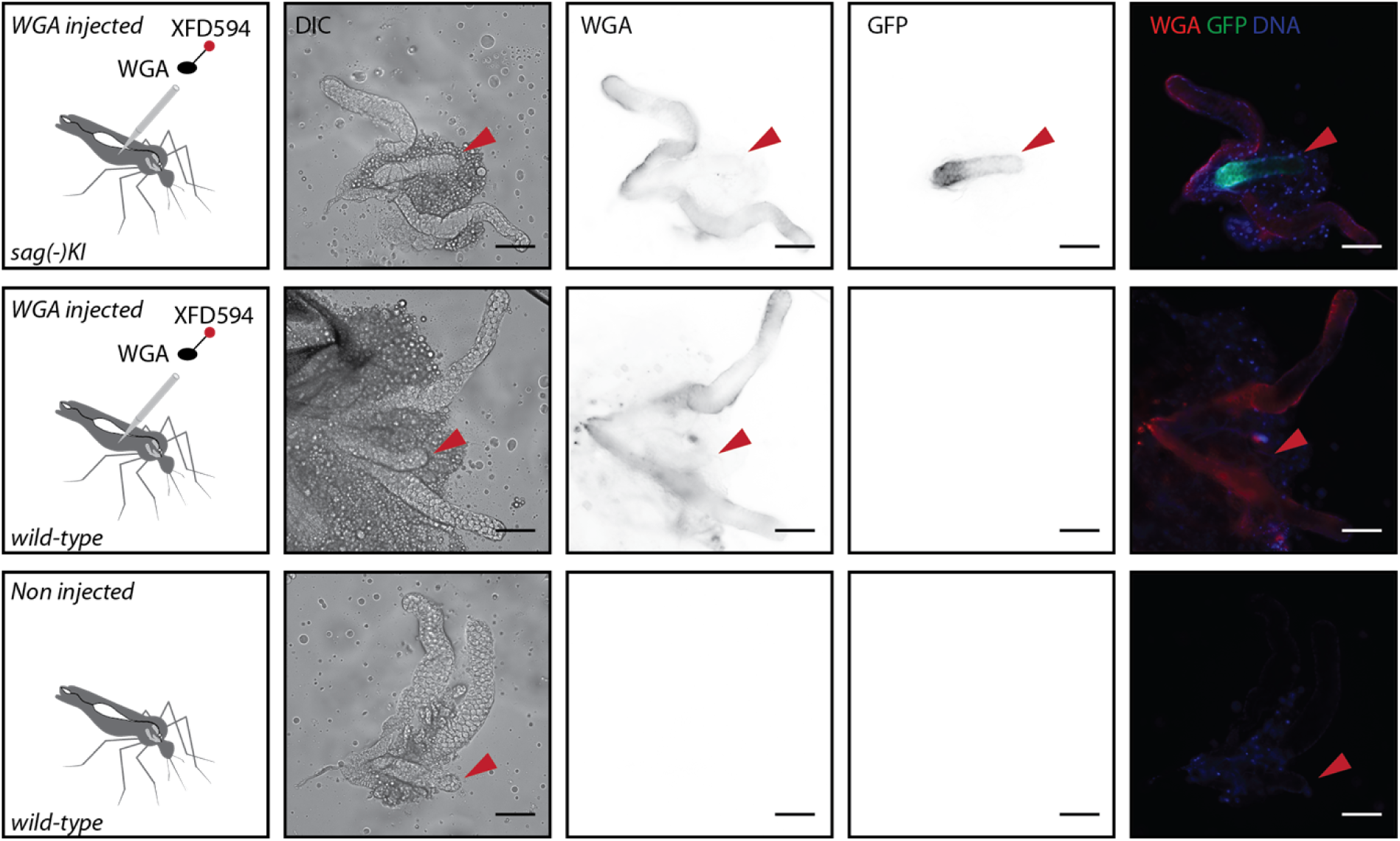
WGA coating is specific upon injection and not affected by the absence of Saglin. Staining patterns of salivary glands dissected from female mosquitoes injected with wheat germ agglutinin (WGA) coupled to XFD594. The salivary gland staining of an injected female homozygous for *sag(-)KI* is compared to two salivary glands dissected from an injected and a non-injected wild-type (*control*) female. Scale bar: 100 µm. Columns from left to right: genotype of injected mosquitoes, differential interference contrast (DIC), WGA and GFP signal in black on white; merge of WGA, GFP and nucleic acid staining (DNA). Red arrows indicate the median lobe.

**Figure 8 – figure supplement 1.**
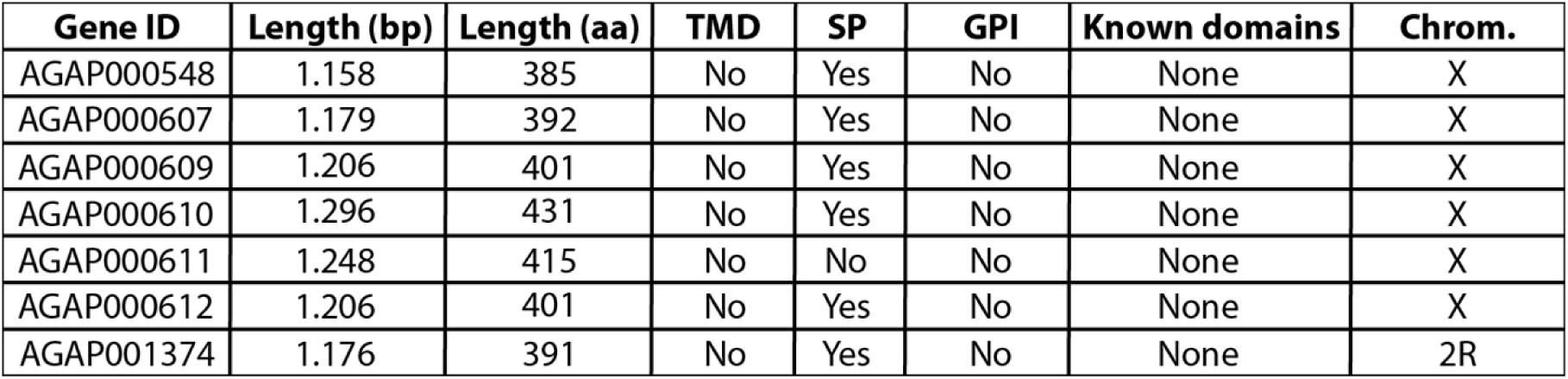
Features of SG1 family proteins. Table summarizing different parameters of all members of the SG1 family as well as predicted transmembrane domains (TMD), signal peptides (SP), GPI-anchors (GPI) and other known domains. Sequence information was retrieved from VectorBase (version 54) (Amos *et al*., 2022). Signal peptide predictions were performed using SignalP 5.0 (Almagro Armenteros *et al*., 2019), TMDs were predicted with TMHMM 2.0 (Krogh *et al*., 2001), GPI-anchors with GPIpred (Pierleoni, Martelli and Casadio, 2008) and other known domains with SMART (Letunic, Khedkar and Bork, 2021).

